# Dynamic zonal identity is a functional determinant of liver disease

**DOI:** 10.64898/2026.11.10.765616

**Authors:** Tyler M. Yasaka, Hiroki Inada, Chang Kyung Kim, Anya Singh-Varma, Shikai Hu, Brandon M. Lehrich, Leon Min, Minakshi Poddar, Anisha Jain, Vik Meadows, Ravi P. Rai, Ian Sipula, Fiona M. Bello, Dillon C. Gavlock, Yasuhito Tanaka, Amber M. Vandevender, Sucha Singh, Yumeng Ren, Fu-Ying Qin, Shikhar Uttam, Aaron Bell, Xiaochao Ma, Reben Raeman, Jia-Jun Liu, Silvia Liu, Jacquelyn A Brown, Mark T. Miedel, Michael J. Jurczak, Yu-Chiao Chiu, Stephane Angers, Nicholas K.H. Khoo, Satdarshan P. Monga

**Author notes:** Corresponding Authors: Satdarshan P. Monga, M.D., Endowed Chair, Professor and Chairman, Department of Pharmacology and Chemical Biology, Director, Organ Pathobiology and Therapeutics Institute, and Pittsburgh Liver Research Center, University of Pittsburgh School of Medicine, 3550 Terrace Street, Scaife Hall, Room A919, Pittsburgh, PA 15213, Nicholas K.H. Khoo, Ph.D. Assistant Professor, Department of Pharmacology and Chemical Biology, Organ Pathobiology and Therapeutics Institute, 3550 Terrace Street, Scaife Hall, Room S927.1, Pittsburgh, PA 15213. These authors contributed equally to this work (Tyler M. Yasaka, Hiroki Inada, Chang Kyung Kim).

## Abstract

Liver zonation demarcates functional division of labor within the hepatic lobule, yet how zonal identity is reshaped during disease remains poorly understood. We developed RIZLiver, a Bayesian framework that infers zonal identity from gene expression without spatial coordinates. RIZLiver identified progressive loss of zone-3 metabolic identity with advancing stages of metabolic dysfunction–associated steatotic liver disease (MASLD) across independent patient cohorts. Complementary mouse models reveal that perturbing zonal identity shifts susceptibility to diet-induced metabolic injury, linking zone-3 identity to adaptive lipid utilization through β-catenin and PPARα. *CTNNB1*-mutated hepatocellular carcinomas (HCC) showed an enriched zone-3 and lipid-oxidative phenotype, extending metabolic perturbations beyond MASLD. Pharmacologic reactivation of the zone-3 program shows a therapeutic effect in MASLD. Conversely, interfering with lipid-metabolic addiction in *CTNNB1*-mutated HCC by targeting PPARα yielded a therapeutic benefit. Our findings identify zonation as a dynamic component of disease phenotype and its restoration as a potential therapy.

**Highlights:**

- RIZLiver infers hepatic zonal identity from gene expression
- MASLD progression is characterized by exhaustion of zone-3 identity
- Wnt-active HCCs acquire a targetable zone-3 reprogramming
- Zone 3 engages a targetable, adaptive lipid utilization program

**Graphical Abstract:** 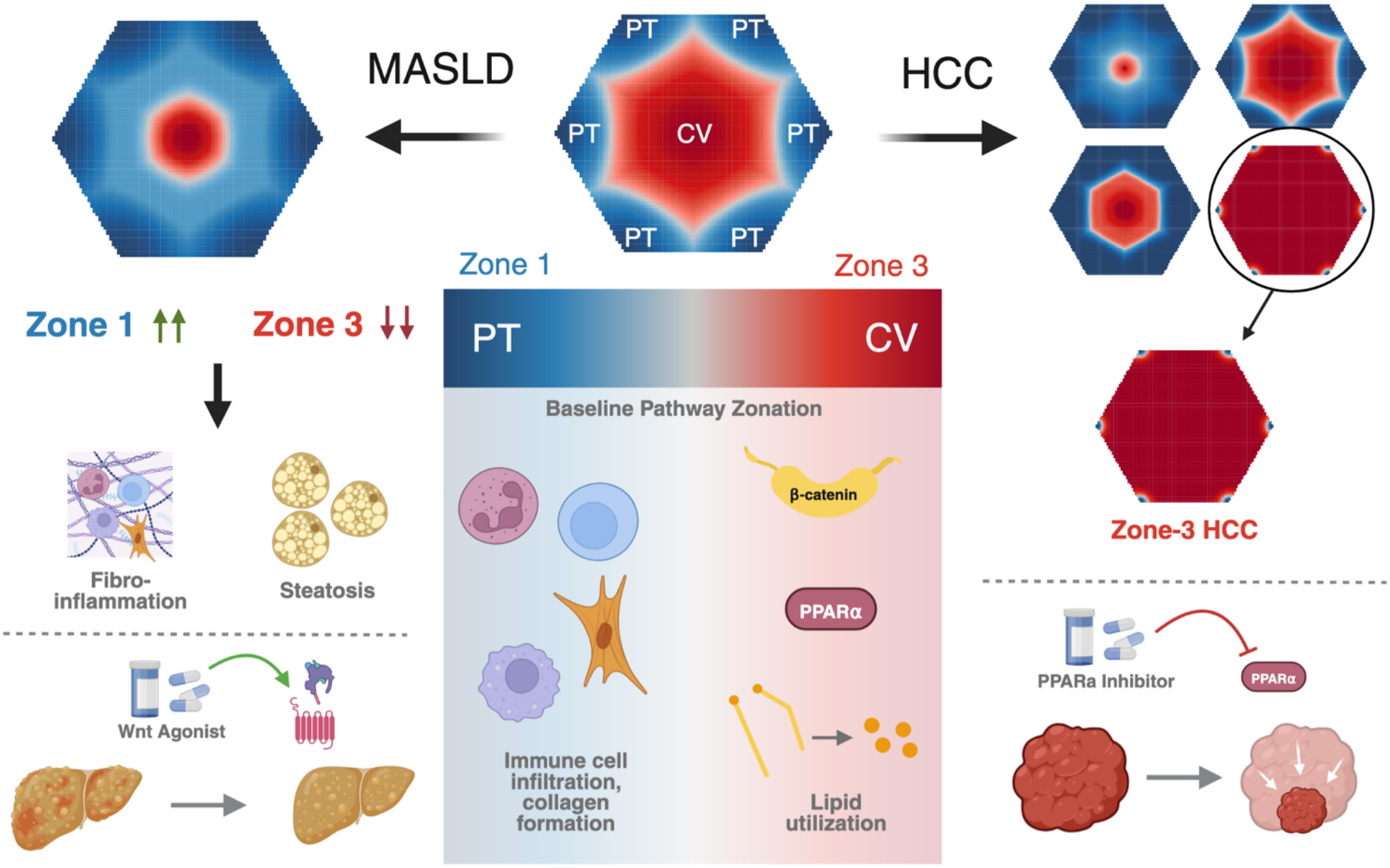

## Introduction

Hepatic zonation is a defining feature of liver organization that dictates cellular function across the liver lobule, spanning the portal triad (zone 1) to the central vein (zone 3), with an ambiguous intermediate zone 2. This spatial arrangement enables hepatocytes to compartmentalize complementary metabolic processes^1^. While zone 1 specializes in functions such as gluconeogenesis and ureagenesis, zone 3 has an established role in glycolysis, xenobiotic metabolism, and bile acid synthesis. Zone 3 has also been associated with de novo lipogenesis, though the capacity of pericentral hepatocytes for lipid utilization is less well defined^2^. This functional heterogeneity, which is essential for normal liver homeostasis, is maintained through Wnt/β-catenin signaling^3–5^. Pericentral, endothelial-derived Wnt ligands are well-established promoters of zone-3 identity, driving β-catenin nuclear translocation and expression of target gene programs, with resulting consequences in metabolic function^6^. Hepatic stellate cells (HSCs) also reside throughout the lobule and have recently been shown to be the source of R-Spondin-3, a well-known enhancer of the Wnt signaling and a contributor of the β-catenin-dependent zone-3 gene expression^7^. While HSCs have been shown to express some Wnt ligands, their contribution to zonal identity is less clear^7,8^.

Multiple liver pathologies, including cholangiopathies, hepatitis, and drug-induced liver injury, demonstrate zonated phenotypes on histological analysis. In adult metabolic dysfunction-associated steatotic liver disease (MASLD), steatosis tends to originate in zone 3, eventually extending to zone 1 with disease progression to metabolic dysfunction-associated steatohepatitis (MASH)^9^. MASLD affects approximately 30% of the global population and carries an increased risk for the development of cirrhosis and hepatocellular carcinoma (HCC)^10^. Recent work in mouse models has proposed that HCC may arise more likely from zone-3 hepatocytes^11^. However, a separate question from zonal origin is whether zonal identity is itself remodeled in liver diseases. Advances in spatial transcriptomics, single-cell technologies, and high-resolution tissue imaging have revealed changes in zonal gene expression, metabolic compartmentalization, and hepatocyte identity in both experimental models and clinical datasets of various pathologies^1,12^. However, whether these alterations are consequences due to adaptation or maladaptation because of the ongoing liver injury and thus contribute to disease susceptibility, remains unclear.

Further, defining the significance of zonation remodeling in human disease has been limited by the difficulty of precisely assigning spatial identity across heterogeneous transcriptomic datasets. While spatial transcriptomic technologies provide direct positional information, assignment of zonal identity remains inconsistent across studies. Conversely, conventional single-cell RNA sequencing (scRNA-seq) and single-nucleus RNA sequencing (snRNA-seq) datasets provide extensive molecular information without spatial coordinates, often relegating zonation characterization to coarse labeling based on a few zonation markers. We therefore developed RIZLiver, a Bayesian inference framework that infers hepatic zonal identity from gene expression, enabling precise delineation of zonal distribution across samples. This approach provides a way to interrogate hepatic zonation across experimental models and disease states, supporting the characterization of zonal identity in human disease along with functional investigation in animal models of perturbed zonation.

Further, we integrated RIZLiver-based analysis of independent human liver disease cohorts with complementary genetic and pharmacological mouse models to determine the balance of hepatic zonal identity in both MASLD and HCC and elucidate novel role of zonated functions in disease trajectories. We identify an initial expansion followed by exhaustion of zone 3-associated metabolic programs as a feature of human MASLD progression and use complementary mouse models that disrupt or expand pericentral hepatocyte identity to determine the functional consequences of this remodeling which were then exploited for novel therapeutic intervention. We further identify a lipid utilization program associated with hepatocyte β-catenin activity that contributes to adaptation to metabolic stress. We find that amid heterogeneous zonal phenotypes in HCC, a zone-3-enriched subset is also metabolically dependent on this program. Finally, we provide evidence that zone-3 program can be pharmacologically modulated for therapy during disrupted zonal identity.

## Results

### RIZLiver enables robust inference of hepatic zonation across platforms and disease states

Computational approaches struggle to precisely and consistently quantify zonation across diverse sequencing platforms, limiting the ability to determine how spatial liver organization changes during disease progression. To address this, we developed RIZLiver, a Bayesian inference algorithm that infers zonal identity from gene expression of cells or subcellular spots across the lobule in a species-specific manner. Importantly, this inference is performed without the use of spatial coordinates, allowing it to be applied to single-cell and single-nucleus datasets. Once baseline zonation has been inferred in the user-supplied data, RIZLiver supports further inference of zonal identity in comparison samples. Comparison samples are assessed relative to the baseline samples, supporting precise quantification and visualization of changes in zonation across physiologic conditions and animal models (**Methods**; **Fig. 1a**).

**Figure 1.**
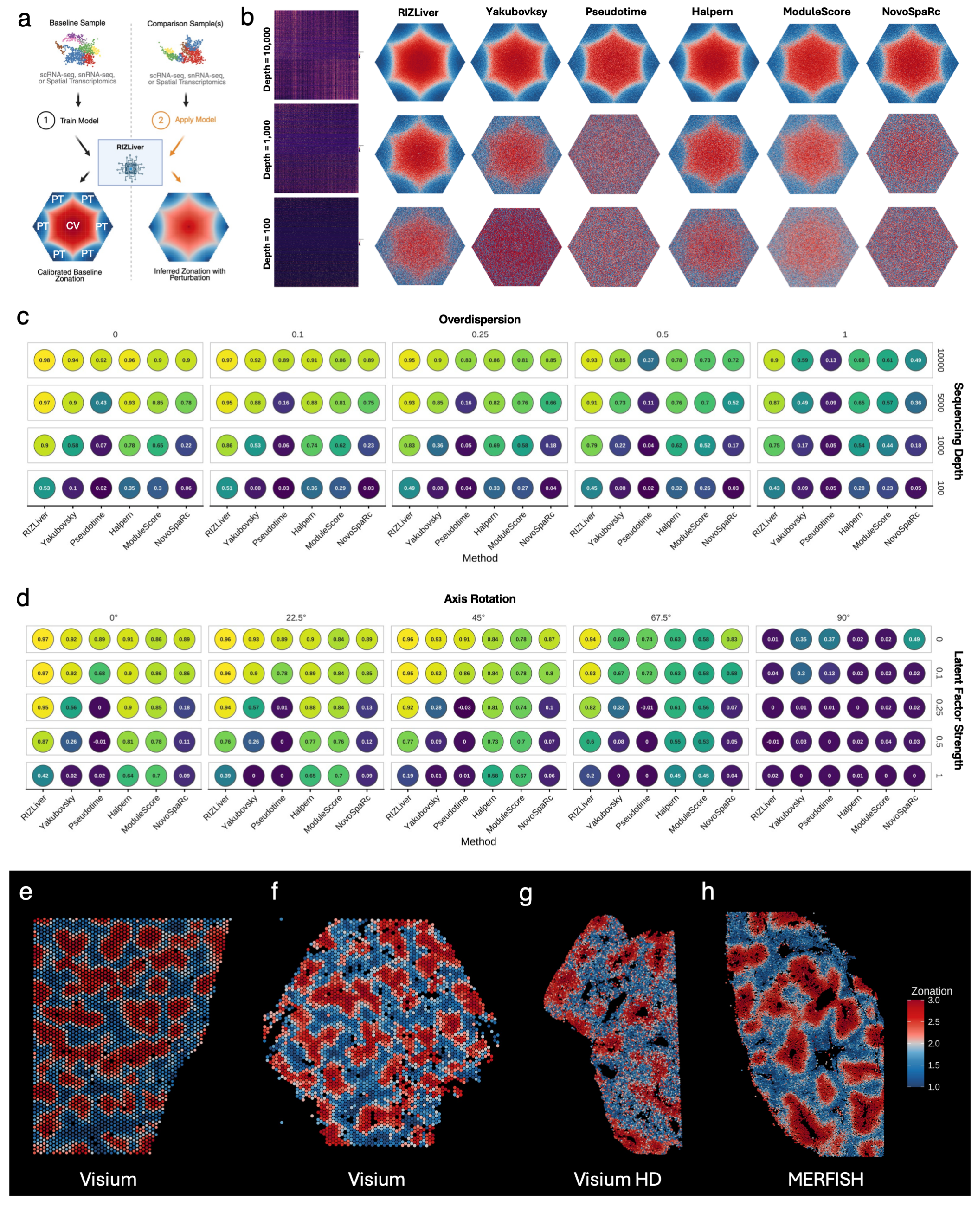
RIZLiver enables robust inference of hepatic zonation. **(a)** Schematic of RIZLiver usage. The user provides a baseline sample for training a dataset-specific model, which can then be applied to additional comparison samples. **(b)** Comparison of zonation inference methods at multiple sequencing depths in simulated experiments. Virtual lobule plots indicate the true location based on the location within the hexagonal lobule representation, with inferred location indicated by color, for one randomly selected experiment for method and sequencing depth. **(c)** Evaluation of zonation inference methods in simulated human liver experiments across multiple biological variability (overdispersion) and technical variability (sequencing depth) values. Displayed values are average Spearman correlation coefficients across 10 simulations. **(d)** Evaluation of zonation inference methods in simulated human liver experiments across multiple axis rotation degrees and latent factor strengths. Displayed values are average Spearman correlation coefficients across 10 simulations. **(e–h)** Visualization of RIZLiver-inferred zonation assignments across four human liver samples spanning three datasets and platforms (Visium, Visium HD, and MERFISH).

To evaluate the ability of RIZLiver to recover baseline zonation from sparse and noisy data, we generated simulated hepatic gene expression data in which each cell’s zonal position was known and evaluated RIZLiver alongside previously utilized zonation inference methods at varying levels of sequencing depth (technical noise) and overdispersion (biological noise; **Methods**). Relative to other methods, RIZLiver consistently achieved superior correlation with the ground truth zonal identity in both simulated human (**Fig. 1b,c, Fig.S1**) and mouse (**Fig.S2a,b**) liver gene expression data.

We next tested robustness to systematic corruption by modeling discrepancy between the reference and the target (axis rotation) and the presence of a dominant external signal (a latent factor). We reasoned that methods which prioritized trust in a predefined reference (e.g. Halpern et al.^13^ and Seurat’s *AddModuleScore* function) would be robust to latent factors but sensitive to axis rotation, while methods which prioritized trust in the target data (e.g. Yakubovsky et al.^14^ and Pseudotime) would be sensitive to latent factors but robust to axis rotation. Our simulations confirmed this tradeoff (**Fig. 1d, Fig.S3, Fig.S4a,b**). While RIZLiver was outperformed at extreme levels of these parameters, it consistently achieved superior performance at more modest values (including combinations of axis rotation and latent factor strength), demonstrating its ability to balance trust in the reference and trust in the target data, achieved using Bayesian inference which treats reference data as the prior and target data as new evidence.

Finally, we benchmarked RIZLiver on manually annotated human spatial transcriptomic data spanning three technologies (Visium, Visium HD, and MERFISH), using datasets independent of the reference. RIZLiver achieved performance either superior to other methods or, in one case, comparable to the highest performing method as evaluated by Spearman correlation and RMSE (**Fig. 1e–h**, **Fig.S5**). In summary, our framework consistently inferred zonal identity from gene expression across wide-ranging conditions in both simulated and real-world evaluations, establishing RIZLiver as a robust and platform-agnostic framework for delineation of hepatic zonation in both preclinical murine models and human liver disease.

### Exhaustion of zone-3 identity is a hallmark of MASLD progression

To establish the zonal orientation of gene expression programs at baseline, we first applied RIZLiver to publicly available Visium spatial transcriptomic data of healthy human liver. Single-sample gene set enrichment analysis (ss-GSEA) of inferred zonation demonstrated trends broadly consistent with established paradigms. Periportal (zone-1) hepatocytes were enriched for extracellular-matrix, complement, and related programs, whereas pericentral (zone-3) hepatocytes were enriched for xenobiotic, steroid, bile-acid, and fatty-acid metabolism, including fatty-acid β-oxidation (**Fig. 2a–f, Fig.S6a,b**).

**Figure 2.**
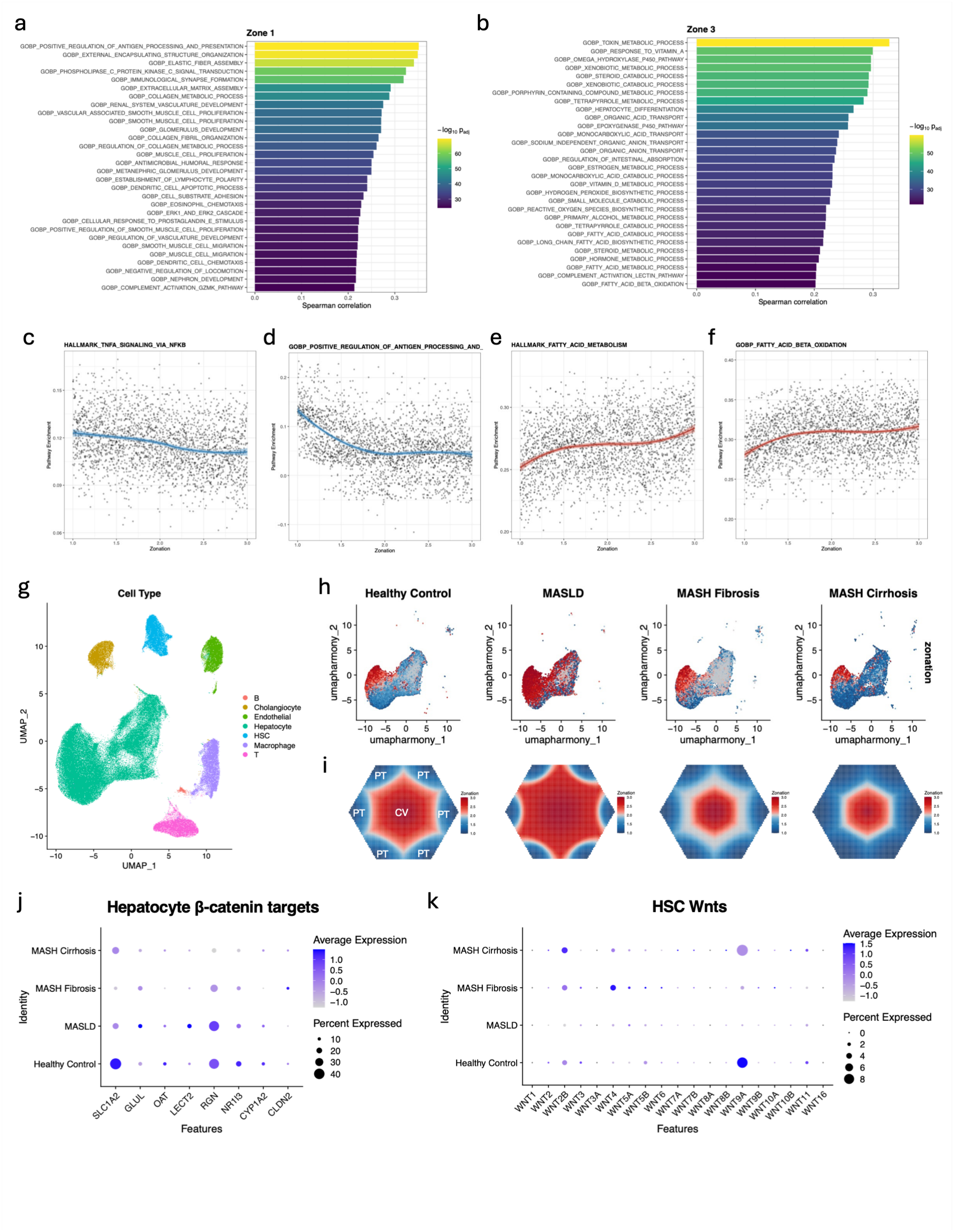
RIZLiver characterizes zonation in homeostasis and throughout MASLD progression. **(a,b)** Top zonated Gene Ontology biological process (GOBP) terms as ranked by Spearman correlation between inferred zonation and single-sample gene set enrichment analysis (ss-GSEA) in Visium spatial transcriptomics of healthy human liver. **(a)** Top zone-1-enriched pathways. **(b)** Top zone-3-enriched pathways. **(c–f)** Scatter plots of selected pathways among the top-ranked hallmark and GOBP terms in zone 1 and zone 3. The x-axis captures the inferred zonation coordinate as a continuous value from the portal triad (1.0) to the central vein (3.0), while the y-axis indicates the ss-GSEA score. **(g)** Uniform Manifold Approximation and Projection (UMAP) visualization of a public snRNA-seq dataset including patients across multiple stages of MASLD. **(h,i)** RIZLiver-inferred zonation score of hepatocytes in snRNA-seq across stages of MASLD projected in **(h)** UMAP space and **(i)** in virtual lobule representations. **(j)** Hepatocyte β-Catenin target gene expression in snRNA-seq across stages of MASLD. **(k)** Total expression of Wnt genes in hepatic stellate cell (HSC) snRNA-seq across stages of MASLD.

The observation that both metabolic and inflammatory pathways were zonated in baseline livers prompted us to ask whether zonation may be altered in MASLD and MASH, which begins with metabolic dysfunction and eventually leads to progressive inflammation and fibrosis. We hypothesized that MASLD progression would not simply reflect uniform transcriptional changes across all hepatocytes but instead would involve progressive and modular remodeling of hepatocyte spatial identity.

To assess this, we first applied RIZLiver to a previously published snRNA-seq dataset (GSE256308)^7^, assessing hepatocyte zonation changes from healthy control livers through early MASLD (F0), MASH fibrosis (F2-F3), and MASH cirrhosis (F4) (**Fig. 2g–i**). This analysis demonstrated an initial zone-3 expansion in MASLD, followed by contraction in MASH. Evaluation of zonal identity scores (**Methods**) relative to baseline further revealed a perturbed zone-3 profile across MASLD stages (**Fig.S6c**). To determine whether this remodeling was conserved across independent cohorts and sequencing approaches, we further applied RIZLiver to a second snRNA-seq dataset comparing healthy controls to F2 and F3 MASH (GSE212837)^15^ (**Fig.S7a,b**), as well as spatial transcriptomic dataset from liver biopsies spanning fibrosis stages in MASLD^16^ (**Fig.S7c,d**). These analyses reproduced the overall finding of progressive diminution of the zone-3 identity in MASH compared to either healthy controls or early MASLD. Taken together, these replicated findings across cohorts and platforms suggest zone-3 expansion as an initial adaptation to metabolic injury which becomes exhausted after persistent and progressive metabolic stress.

Given the central role of the canonical Wnt/β-catenin signaling in maintaining zone-3 identity^1^, we asked whether zone 3 contraction in MASLD is accounted for by decreased overall abundance of Wnt ligands. However, analysis of previously published human snRNA-seq^7^ revealed a disconnect between Wnt ligand expression and downstream canonical β-catenin signaling. Hepatocyte expression of canonical β-catenin target genes, including *GLUL*, *OAT, CYP1A2, LECT2*, and *RGN*, broadly mirrored the trajectory of inferred zone-3 identity with initial increase over baseline in early MASLD followed by loss during MASH fibrosis and cirrhosis (**Fig. 2j**). However, endothelial cells (ECs), which promote canonical β-catenin signaling in hepatocytes via *WNT2* and *WNT9B*^6^, showed an initial increase in expression of these two ligands followed up to MASH fibrosis followed by notable decrease in MASH cirrhosis (**Fig.S7e**). This decoupling suggests that altered endothelial Wnt abundance alone does not account for the progressive loss of hepatocyte pericentral identity. To more holistically address Wnt gene expression in MASLD, we looked at the expression of all Wnts in MASH liver dataset. Overall, expression of several Wnt ligands remained preserved or even paradoxically increased during MASH progression (**Fig.S7f).** This was especially true of *WNT4*, *WNT5A*, *WNT3* and *WNT2B*, which have independently been shown to negatively impact β-catenin activity in various contexts^17–20^. Given that progressive MASH is associated with HSC activation and increasing fibrosis, we queried for alterations in Wnt expression in HSCs. Intriguingly, we observed increases in HSC expression of *WNT2B*, *WNT4*, and *WNT5A* during MASH fibrosis and/or cirrhosis over baseline **(Fig. 2k**). Collectively, these observations suggest that progressive loss of the zone-3 transcriptional program may not be explained simply by diminished endothelial cells canonical Wnt abundance, except during MASH cirrhosis where their absence is unequivocal. Further, it raises the possibility that some non-canonical Wnts from HSCs may be contributing to loss of hepatocyte spatial identity and function.

### HSC-derived Wnt secretion remodels hepatic zonation

Having observed an increase in HSC-derived, non-canonical Wnt expression coinciding with a decline in hepatocyte β-catenin activation, diminished zone-3 identity, and MASLD progression, we posited that HSC-derived Wnts may be contributing to MASH pathogenesis. Using a model of HSC-specific depletion of Wnt secretion, achieved by crossing lecithin retinol acyltransferase (*Lrat*)-Cre with *Wls*^flox/flox^ mice (HSC-Wls-KO, **Fig. 3a**), we had previously reported HSC-derived Wnts to be dispensable for zone-3 gene expression of *Glul*^8^. However, to address this more holistically, we performed snRNA-seq on HSC-Wls-KO livers and assessed metabolic zonation using RIZLiver. Intriguingly, we observed a notable expansion in zone-3 identity in these livers at the expense of zone 1 (**Fig. 3b,c, Fig.S8a**) without any evidence of HSC activation (**Fig.S8b**). These observations were validated by immunohistochemistry (IHC) that was performed for multiple zonation markers (**Fig. 3d,e**). The basis for this expansion is unclear but likely reflects a culmination of impaired secretion of β-catenin inhibitory Wnts from HSCs along with compensatory shifts within the hepatic Wnt network such as increased expression of *Wnt2* in HSC-Wls-KO ECs (**Fig. 3f**). Incidentally, HSC-Wls-KO provides us with a unique *in vivo* model of expanded zone-3, enabling us to directly address the role of the zone-3 module in hepatic pathobiology.

**Figure 3.**
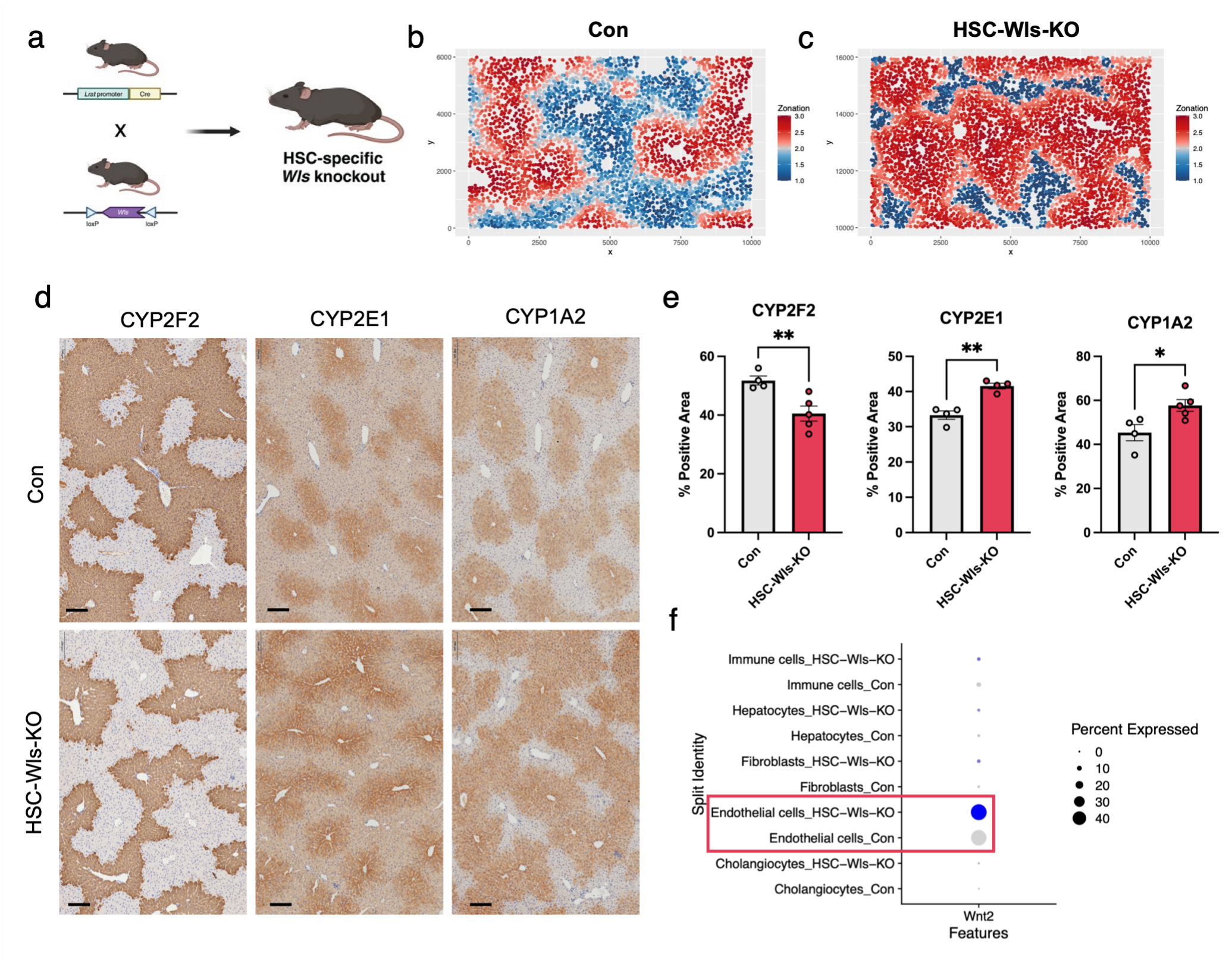
Depletion of HSC-derived Wnts remodels hepatic zonation. **(a)** Schematic illustrating the generation of hepatic stellate cell (HSC)-specific *Wls* knockout mice by crossing Lrat-Cre mice with *Wls*^fl/fl^ mice. **(b, c)** Spatial plots of inferred zonation scores using RIZLiver in HSC-Wls-KO and littermate floxed controls (Con). The zonation score indicates inferred position as a continuous score from the portal triad (1.0) to the central vein (3.0). **(d)** Representative immunohistochemical (IHC) staining of zonation markers in HSC-Wls-KO and control. **(e)** Quantification of IHC of zonation markers in HSC-Wls-KO and control. **(f)** Dot plot of Resolve spatial transcriptomics *Wnt2* expression across cell types between control and HSC-Wls-KO. For (e), data are presented as mean ± SD; two-tailed Student’s t-test was used for statistical analysis, with significance denoted by p < 0.05 (*) and p < 0.01 (**).

### Hepatic zonation remodeling is associated with altered metabolic function

Investigational snRNA-seq of HSC-Wls-KO hepatocytes ranked zone-3 enriched processes – fatty acid, bile acid, and xenobiotic metabolism – among the most enriched pathways (**Fig.S9a**). We therefore asked whether HSC-Wls-KO exhibit concordant metabolic functions. To directly assess hepatic mitochondrial function, high-resolution respirometry on mitochondrial-enriched liver lysates was performed. This revealed no genotype difference in basal respiration or ADP-stimulated oxidative phosphorylation on any substrate, indicating comparable baseline or coupling efficiency of mitochondrial respiration (**Fig. 4a,b**). Genotype-dependent differences emerged only under increased respiratory demand. Succinate stimulation (Complex II) and FCCP-induced uncoupling both produced significantly greater responses in HSC-Wls-KO mitochondria; this was observed under lipid-supported (palmitoyl-carnitine/malate) but not carbohydrate-supported (pyruvate/malate) conditions. Loss of HSC-derived Wnt secretion therefore does not globally increase mitochondrial activity but selectively enhances lipid-oxidative capacity under metabolic demand. Consistent with expanded zone-3 metabolic function, HSC-Wls-KO livers also exhibited an increased total bile-acid pool (**Fig.S9b**) without any changes in its overall composition (**Fig.S9c,d**).

This hepatic remodeling was accompanied by a favorable whole-body metabolic phenotype. HSC-Wls-KO mice were leaner than littermate floxed controls, with decreased fat mass and increased lean mass by ¹H-NMR (**Fig. 4c,d**, **Fig.S9e**. Food intake was unchanged, and HSC-Wls-KO mice exhibited reduced physical activity (**Fig. 4e**, **Fig.S9f**). Genotype-dependent differences in energy expenditure (EE) were not observed after accounting for lean mass (**Fig. 4f**, **Fig. S9g,h**) or body weight (not shown). Therefore, the lean phenotype of HSC-Wls-KO cannot be readily explained by differences in food consumption or locomotor activity.

**Figure 4.**
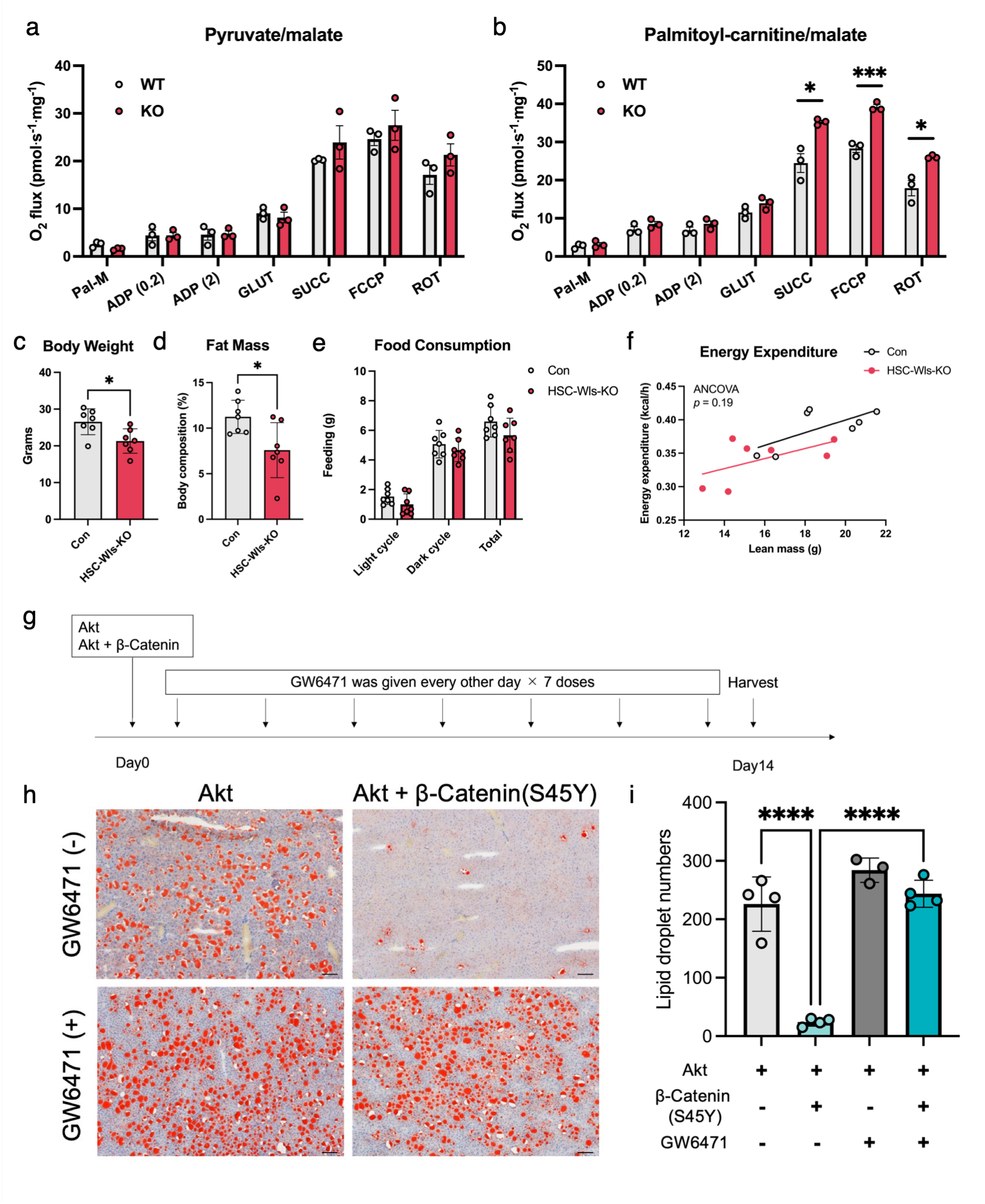
Hepatic zonation remodeling is associated with altered metabolic function. Oxygen consumption rates (O₂ flux) were measured following sequential addition of ADP, glutamate (GLUT), succinate (SUCC), the uncoupler FCCP, and the complex I inhibitor rotenone (ROT). High-resolution respirometry of liver mitochondria using carbohydrate-based substrates (pyruvate/malate) **(a)** and lipid-based substrates (palmitoyl-carnitine/malate) **(b)**. **(c)** Body weight and **(d)** fat mass percentage in control and HSC-Wls-KO mice. **(e)** Food intake (g/kg) during light cycle, dark cycle, and overall. **(f)** Energy expenditure analyzed by ANCOVA with lean mass as a covariate. **(g)** Experimental design for pharmacological inhibition of PPARα using GW6471 in mice expressing Akt or Akt + β-catenin (S45Y). GW6471 was administered every other day for a total of seven doses prior to harvest. **(h)** Representative Oil Red O staining in mouse livers expressing Akt ± S45Y β-catenin and ± GW6471. **(i)** Quantification of lipid droplet numbers. Data are presented as mean ± SD. Statistical analyses were performed using unpaired two-tailed Student’s t-test or one-way ANOVA followed by Tukey’s multiple-comparisons test. Significance is denoted as follows: p < 0.05 (*), p < 0.001 (***), and p < 0.0001 (****).

Having established a correlation between zone-3 expansion and lipid oxidative capacity in HSC-Wls-KO livers, we next sought to probe the hepatocyte-intrinsic mechanism driving this phenotype. PPARα is a major transcriptional regulator of fatty acid oxidation (FAO) and a direct β-catenin target^21,22^ enriched in zone-3^23^ (**Fig.S10a**). We therefore asked whether β-catenin activation was associated with engagement of PPARα-mediated lipid metabolic programs in the setting of steatosis. To test this directly, we applied a more reductionist approach to induce hepatocyte steatosis using hydrodynamic delivery of myristoylated Akt which produces marked lipid-droplet accumulation in cells expressing the exogenous plasmid^24^ (**Fig.S10b,c**). Simultaneous forced co-expression of constitutively active β-catenin (S45Y) in Akt-expressing hepatocytes dramatically reduced steatosis (**Fig.S10d**), demonstrating that hepatocyte β-catenin activation limits lipid accumulation. Consistent with engagement of PPARα-associated lipid utilization, β-catenin activation increased hepatic expression of *Ppara* and its downstream target *Acsl1*, while other FAO genes showed modest or variable changes and lipogenic genes remained largely unchanged (**Fig.S10e**), indicating selective activation of a lipid-oxidation rather than a lipogenic program. To determine whether PPARα was required for this protection, mice were treated with a PPARα antagonist, GW6471, following Akt or Akt + β-catenin (S45Y) delivery (**Fig. 4g**). Pharmacologic inhibition of PPARα largely abolished the anti-steatotic effect of β-catenin activation, restoring lipid accumulation in Akt + β-catenin livers (**Fig. 4h,i**). Together, these gain-of-function experiments demonstrate that hepatocyte β-catenin activation is sufficient to limit steatosis and support contribution of PPARα-associated lipid metabolism to this response.

### Zonal identity bidirectionally determines susceptibility to diet-induced liver injury

Having shown that β-catenin activation is sufficient to limit lipid accumulation through PPARα, we next examined the consequences of losing zone-3 identity. Given the zone-1 preponderance of ECM- and immune-related pathways, we hypothesized that loss of hepatocyte β-catenin would drive zone-1 expansion and result in MASLD disease progression to MASH with steatohepatitis and fibrosis. Chromium Flex snRNA-seq of mouse livers with inducible deletion of *Ctnnb1* in hepatocytes (HC-Ctnnb1-KO, **Fig. 5a**) confirmed a loss of zone-3 identity in favor of zone-1 expansion using the RIZLiver (**Fig. 5b, Fig.S11a,b**). This was validated via IHC (**Fig.S11c**). IPA and CellChat analysis further demonstrated upregulation of immune cell infiltration and pro-fibrotic interactions in HC-Ctnnb1-KO livers (**Fig. 5c**, **Fig.S11d**–**f**), supporting that expansion of zone-1 promotes a pro-inflammatory hepatic milieu.

**Figure 5.**
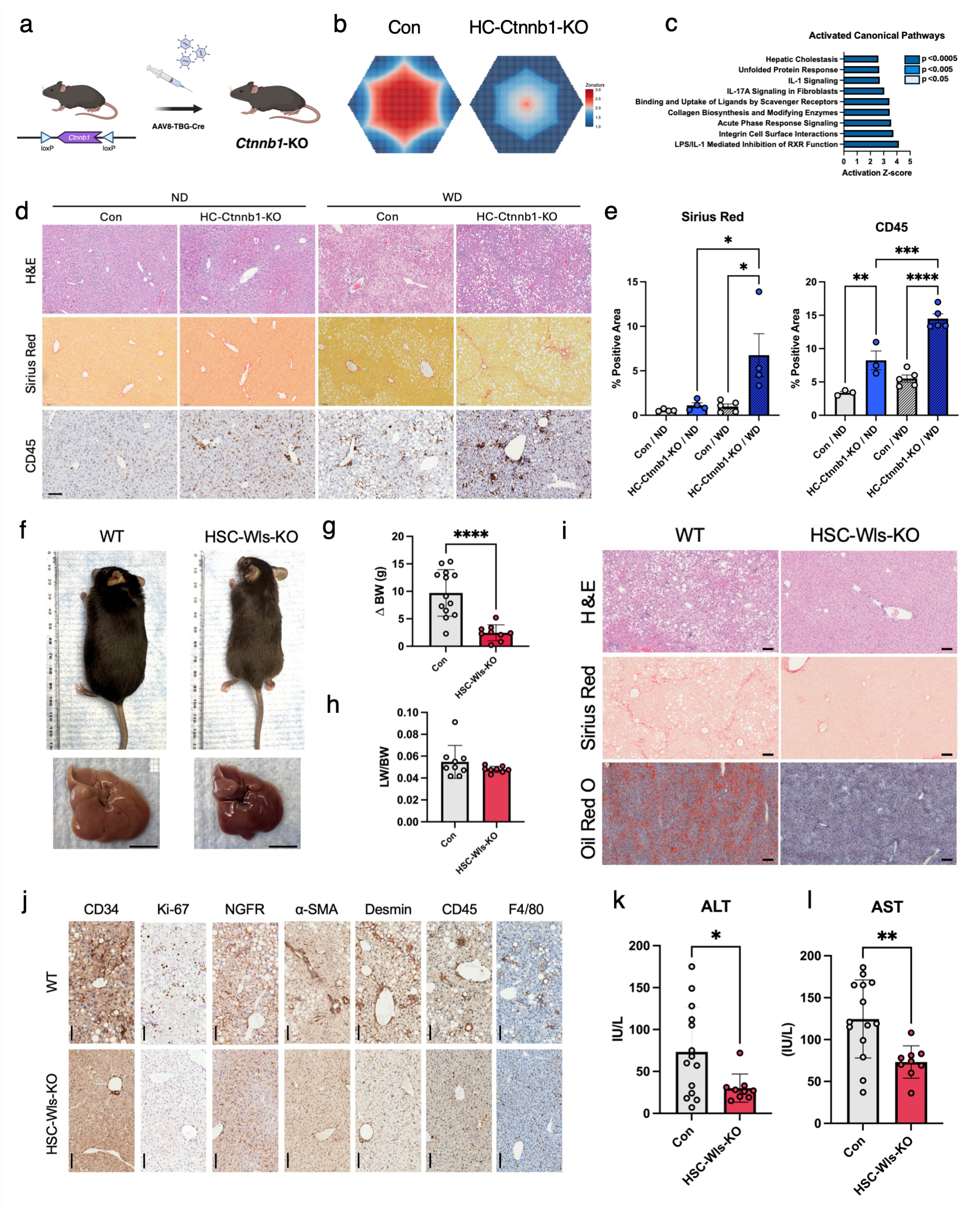
Zonal identity bidirectionally determines susceptibility to diet-induced liver injury. **(a)** Schematic illustrating the generation of inducible hepatocyte (HC)-specific *Ctnnb1* knockout mice by injecting *Ctnnb1*^fl/fl^ mice with AAV8-TBG-Cre. **(b)** Virtual lobule plots of inferred zonal identities using RIZLiver in Chromium Flex snRNA-seq of HC-Ctnnb1-KO and control livers. **(c)** Activated canonical pathways in HC-Ctnnb1-KO mice found by Ingenuity Pathway Analysis (IPA) of Chromium Flex snRNA-seq. **(d)** Representative hematoxylin and eosin (H&E), Sirius red, and IHC staining of immune cells (CD45) in HC-Ctnnb1-KO and control livers ± WD (n = 2 - 5). **(e)** Quantification of % positive area of staining in **(d)**. **(f)** Representative gross images of HSC-Wls-KO mice and livers after 12 weeks of WD. **(g)** Change in body weight (ΔBW) and **(h)** liver weight to body weight ratio in HSC-Wls-KO mice and controls over the course of WD feeding. **(i)** Representative histological images of liver sections stained with H&E, Sirius Red, and Oil Red O in HSC-Wls-KO livers compared with controls following WD. **(j)** Representative IHC staining of liver sections for macrophages (F4/80), immune cells (CD45), hepatic stellate cells (Desmin, α-SMA, Ngfr), proliferation (Ki67), and endothelial cells (CD34) in HSC-Wls-KO and control livers following WD. **(k,l)** Serum biochemistry of **(k)** ALT and **(l)** AST in HSC-Wls-KO versus control livers following WD. Data are presented as mean ± SD. Statistical analyses were performed using unpaired two-tailed Student’s t-test or one-way ANOVA followed by Tukey’s multiple-comparisons test. Significance is denoted as follows: p < 0.05 (*), p < 0.01 (**). p < 0.001 (***), and p < 0.0001 (****).

We then asked whether loss of β-catenin in hepatocytes promotes progression of MASLD to MASH. HC-Ctnnb1-KO mice were placed on normal chow or western diet for 4 weeks, after which β-catenin was deleted, and mice were maintained on diet until week 12 (**Fig.S12a**). This resulted in a significant increase in CD45+ cell accumulation and worsening of fibrosis, suggesting that loss of hepatocyte β-catenin is sufficient to drive progression from steatosis to steatohepatitis (**Fig. 5d,e**).

Conversely, we examined whether zone 3 expansion is protective against metabolic injury by subjecting HSC-Wls-KO mice to WD for 12 weeks. Macroscopic examination revealed a visibly leaner phenotype and reduced liver steatosis in KO mice (**Fig. 5f**), with reduced weight but unchanged liver weight-to-body weight ratios (LW/BW; **Fig. 5g,h**), indicating that improved hepatic pathology was not attributable to differences in liver size. In contrast, visceral adipose tissue (VAT)-to-body weight ratios (VAT/BW) were significantly reduced in KO mice (**Fig.S12b**), consistent with the lean phenotype and enhanced lipid utilization observed in these animals. Food intake was comparable between groups (**Fig.S12c**), suggesting that improved liver outcomes were not due to altered caloric intake.

Histologically, HSC-Wls-KO livers showed marked attenuation of fibrosis and lipid accumulation by H&E, Sirius Red, and Oil Red O (**Fig. 5i**). Inflammation and fibrogenesis were broadly suppressed, with decreased staining for F4/80, CD45, Desmin, α-SMA, and NGFR and reduced hepatocyte proliferation assessed by Ki-67 (**Fig. 5j**), accompanied by downregulation of *Tgfb1*, *Ccl2*, *Tnf*, *Ptprc*, and *Col1a1* (**Fig.S12d**). Consistent with these findings, serum ALT and AST were significantly reduced in HSC-Wls-KO mice (**Fig. 5k,l**), while cholesterol, triglyceride, and glucose levels showed a trend toward reduced levels without reaching statistical significance in the non-fasted state (**Fig.S12e**). Together, these findings indicate broad protection from MASLD pathology in HSC-Wls-KOs.

Collectively, complementary gain- and loss-of-function models demonstrate that hepatic zonation is not simply a passive marker of metabolic disease progression but actively influences adaptation to WD-induced metabolic stress. Expansion of zone-3 identity through HSC-Wls deletion promotes a metabolically adaptive state associated with preferential fatty acid (FA) substrate utilization and resistance to WD-induced liver injury, whereas loss of hepatocyte β-catenin disrupts zone-3 programs and increases inflammation and fibrotic responses. These results suggest that MASLD progression may reflect not simply altered hepatic zonation, but failure of adaptive zonation remodeling required to maintain metabolic liver homeostasis during chronic nutrient and metabolic stress.

### Zone-3 metabolic programming is retained in β-catenin-driven HCC

Our preceding studies indicate that hepatic zonation represents a dynamic metabolic program that influences adaptation to liver injury. Expansion of zone-3 programming enhances hepatic lipid utilization and protects against WD-induced liver injury, whereas loss of pericentral identity increases disease susceptibility. Adaptive biological programs, however, may become pathological when constitutively activated. We therefore asked whether the same zonation-associated metabolic programs that promote hepatocyte adaptation are retained and leveraged in HCC, of which a substantial fraction is driven by constitutive β-catenin activation^25^.

To characterize hepatic zonation in HCC, we profiled six human HCCs across disease stages along with a non-tumor control liver using the Xenium spatial transcriptomic platform (**Table S1**). RIZLiver analysis revealed both intra- and inter-tumoral heterogeneity in zonation phenotypes (**Fig. 6a,b, Fig.S13a**). Tumors differed in both the distribution and overall strength of these identities, with broad weakening of zonal identity consistent with malignant transformation (**Fig.S13b**), indicating that zonation phenotypes are variably preserved across HCCs.

**Figure 6.**
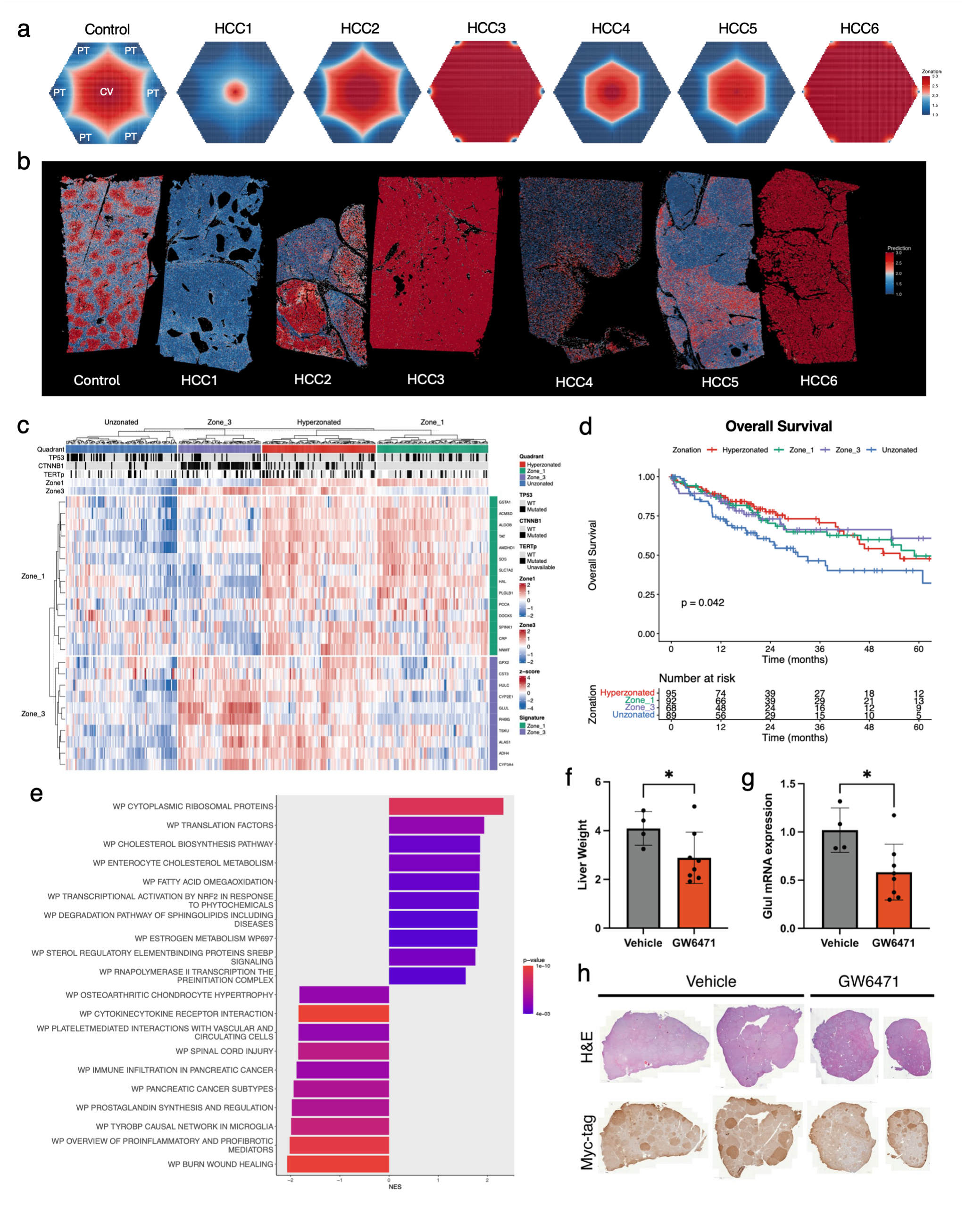
RIZLiver characterizes zonation phenotypes in hepatocellular carcinoma. **(a)** Virtual lobule plots of inferred zonation across 1,354,381 cells in a healthy control liver and six hepatocellular carcinoma samples captured using Xenium spatial transcriptomics. A RIZLiver model was trained on the control sample and applied to the tumor samples. **(b)** Spatial plots of inferred zonation scores from **(a)**. **(c)** Zonation subtypes of HCC in The Cancer Genome Atlas (TCGA) derived from HCC-specific zonation signatures generated from the Xenium dataset. Zone-1 and zone-3 signatures were used to label samples as *unzonated*, *zone-1*, *zone-3*, or *hyperzonated*. *TP53*, *CTNNB1*, and *TERT* promoter mutation status, along with zone-1 and zone-3 signatures, are indicated along with expression of the signature genes. **(d)** Kaplan-Meier plot depicting overall survival of HCC patients in TCGA-LIHC stratified by zonation subtype. Log-rank p-value as indicated. **(e)** Gene set enrichment analysis (GSEA) of up- and down-regulated pathways in zone-3 HCC. **(f)** Liver weight and **(g)** Glul mRNA expression in the *CTNNB1-NFE2L2* murine HCC model ± 4 weeks of treatment with GW6471. **(h)** Representative hematoxylin and eosin (H&E) and Myc-tag staining in *CTNNB1-NFE2L2* mice ± 4 weeks of treatment with GW6471. Data are presented as mean ± SD; two-tailed Student’s t-test was used for statistical analysis, with significance denoted by p < 0.05 (*).

To relate zonal phenotype to molecular profiles and outcomes in a larger cohort, we derived HCC-specific zonation signatures from RIZLiver-assigned zonal identities in the Xenium data (**Methods**) and applied them to bulk RNA sequencing in The Cancer Genome Atlas Liver Hepatocellular Carcinoma (TCGA-LIHC) cohort. Clustering by these signatures revealed not only zone-1- and zone-3-oriented subtypes, but also intermediate subtypes denoted as *unzonated* or *hyperzonated* (**Fig. 6c**). The zone-3 signature was increased with *CTNNB1* and *TERT* promoter mutations and negatively associated with *TP53*, whereas the zone-1 signature was reduced in *TP53* and *CTNNB1* mutated tumors (**Fig.S13c**), supporting that oncogenic β-catenin activation preserves, rather than erases, features of the normal zone-3 program. Zonation subtypes further differentiated patient outcomes across multiple survival metrics, with the unzonated group demonstrating the poorest survival (**Fig. 6d, Fig.S13d,e**). These findings are consistent with prior work which found that periportal- and pericentral-type HCCs are associated with improved survival outcomes, with *CTNNB1* mutation associated with pericentral-type HCCs and *TP53* mutation associated with a subset of the proliferation class which excludes periportal- and pericentral-type HCCs^26^.

We next asked whether zone-3 HCC carries a targetable metabolic dependency. Gene set enrichment analysis (GSEA) comparing zone-3 tumors with other subtypes ranked fatty-acid ω-oxidation among the most enriched pathways (**Fig. 6e**). Prior work demonstrated that β-catenin-activated tumors depend on FAO and that loss of PPARα attenuates tumor growth^22^. Together with the enrichment of lipid-utilization programs in our zone-3 HCCs, these findings led us to hypothesize that PPARα-dependent lipid utilization may represent a therapeutic vulnerability in zone-3 HCC. Testing this required an *in vivo* model of zone-3 HCC. GSEA of zone-3 tumors revealed enrichment of NRF2 activation (**Fig.S13f**), supporting the *CTNNB1-NFE2L2* HCC model as a zone-3-like HCC model^27^ (**Fig.S13g**). Treatment of established tumors with the PPARα antagonist GW6471 reduced tumor burden, lowering liver weight and *Glul* expression relative to vehicle (**Fig. 6f–h**). Our findings further support previous observations that β-catenin-driven HCC is dependent on PPARα-regulated FAO^22^ and identify a therapeutically exploitable metabolic vulnerability in zone-3 HCC. Thus, features of the lipid-utilization program associated with adaptive zone-3 hepatocyte function during liver injury are retained in β-catenin-driven tumors.

### Pharmacologic Wnt activation engages the zone-3 program to ameliorate steatotic liver disease

We next asked whether conversely, pharmacologic activation of Wnt signaling, could engage zone-3 functions established western diet (WD)-induced liver injury. Our prior work showed that the tetravalent Wnt agonist FL6.13 can restore hepatic zonation and promote liver regeneration in drug induced liver injury model^6^. Because both genetic zone-3 expansion and hepatocyte β-catenin activation limited steatosis, we asked whether pharmacologic Wnt activation could activate zone-3 programs in established WD-induced metabolic injury to have a therapeutic benefit. We administered a tailored tetravalent antibody FL6.13 with a direct Wnt agonist function^6,28^, to mice with established WD-induced disease (dosing from week 12), comparing animals kept on WD (Group A) with animals switched to normal diet (Group B; **Fig. 7a**). FL6.13 reduced body weight within nine days (**Fig. 7b**). Its effect on steatosis, however, was context dependent. In mice maintained on WD, FL6.13 did not significantly reduce hepatic steatosis, whereas in mice switched to normal diet, it produced a marked reduction in lipid accumulation (**Fig. 7c–e**). VAT/BW ratios were unchanged, whereas LW/BW ratios were significantly increased in the FL6.13-treated group (**Fig.S14a**) suggesting that the early systemic effects of FL6.13 were insufficient to overcome the persistent metabolic stress imposed by continued WD feeding and perhaps a more chronic administration of the agent would be needed. This improvement in steatosis was accompanied by an increased proportion of β-catenin target Tbx3-positive zone-3 hepatocytes (**Fig.S14b,c**), consistent with zone-3 hepatocyte reprogramming under metabolic conditions, which appears to be permissive for recovery.

**Figure 7.**
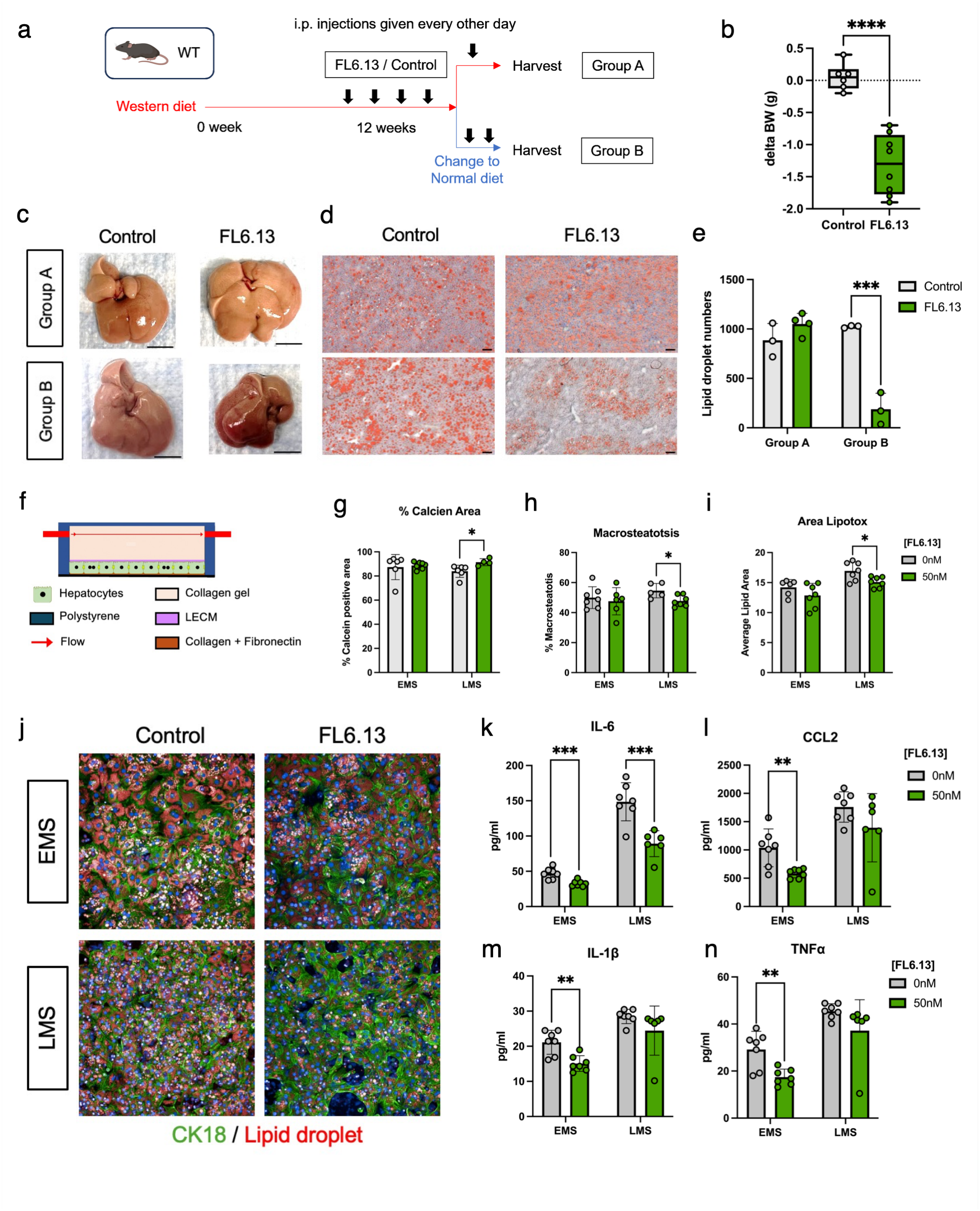
Pharmacologic Wnt activation reactivates the zone-3 program to ameliorate steatotic liver disease. **(a)** Experimental schematic of FL6.13 administration in WT mice subjected to Western diet (WD). After 12 weeks of WD feeding, mice received intraperitoneal injections of FL6.13 or control every other day. Mice were either maintained on WD (Group A) or switched to normal diet (Group B) prior to harvest. **(b)** Change in body weight (ΔBW) at day 9 following treatment with FL6.13. **(c)** Representative gross liver images from Group A and Group B ± FL6.13. **(d)** Representative Oil Red O staining of liver sections from Group A and Group B ± FL6.13. **(e)** Quantification of **(d)**. **(f)** Schematic of the human hepatocyte liver-on-chip system and experimental design. Hepatocytes were cultured under flow conditions and exposed to early metabolic syndrome (EMS) or late metabolic syndrome (LMS) media, followed by treatment with the Wnt agonist FL6.13 or control. **(g–i)** Quantification of **(g)** hepatocyte viability (% Calcein-positive area), **(h)** macrosteatosis (% macrosteatotic cells), and **(i)** lipid accumulation (average lipid area). **(j)** Representative immunofluorescence images of hepatocytes stained for CK18 (green) and lipid droplets (red). **(k–n)** Cytokine profiling of culture supernatants (IL-6, CCL2, IL-1β, and TNFα). Data are presented as mean ± SD. Statistical significance was determined by unpaired two-tailed t-test and denoted as follows: p < 0.05 (*), p < 0.01 (**). p < 0.001 (***), and p < 0.0001 (****).

To determine whether these effects extend to human hepatocytes, we used a hepatocyte-only liver acinus microphysiological system (LAMPS) maintained under perfusion conditions designed to approximate zone-3 oxygen tension and exposed to media modeling key features of MASLD and MASH.^29^ Early metabolic syndrome (EMS; approximating MASLD) medium contained oleic and palmitic acids, elevated glucose and insulin, and reduced glucagon. Late metabolic syndrome (LMS; approximating MASH) medium additionally contained TGF-β1 and lipopolysaccharide (LPS) to introduce profibrotic and inflammatory stimuli, respectively (**Methods**, **Fig. 7f**). Under EMS, FL6.13 significantly reduced secreted inflammatory cytokines measured in the effluent using Luminex including IL-6, IL-1β, CCl2 and TNFα, while it did not significantly alter hepatocyte viability or steatosis scores in the LAMPS **(Fig. 7g-n).** Under LMS, FL6.13 improved hepatocyte viability and reduced macrosteatosis and lipid accumulation, and restricted some, but not all, inflammatory and fibrogenic cytokine levels (**Fig. 7g–n, Fig.S14d**). In summary, in both the mouse and human systems, Wnt activation was beneficial in steatotic liver disease, with the balance of effects differing by context: anti-inflammatory in earlier or otherwise permissive contexts, and anti-steatotic under advanced inflammatory conditions, consistent with a context-dependent restoration of zone-3 function. Together, these findings demonstrate that pharmacologic Wnt activation can engage zone-3 programming in both mouse and human models of steatotic liver disease.

## Discussion

Our findings suggest that hepatic zonal identity is not a static feature of liver architecture but a dynamic ecosystem that is actively remodeled during disease, which impacts how the liver adapts to metabolic stress. Two observations support this view. First, zone-3 identity is exhausted in human MASH, implicating failed zonal remodeling as a feature of progressive metabolic injury. Second, manipulation of zonal identity in mice is sufficient to shift susceptibility to metabolic injury in either direction, establishing that hepatic zonation is functionally relevant in disease outcomes. These observations point to a pericentral metabolic program associated with β-catenin-dependent activity that can support adaptation in the steatotic liver while also being retained in β-catenin-activated HCC. That the same program can be pharmacologically restrained in one disease context and reactivated in another, positions zonation as a dynamic lever governing disease outcomes.

Spatial lipidomic, transcriptomic, and multi-omic studies have identified changes in hepatic lipid distribution and metabolic pathways in human MASLD^30,31^. Building on this work, we demonstrate that zonal identity itself evolves throughout disease progression. Importantly, the contraction of pericentral identity should not be interpreted as uniform loss of every function normally associated with zone 3. Our observations reflect disruption of a coordinated transcriptional state encompassing multiple metabolic programs, which is consistent with the finding that zonated populations become progressively less distinct in MASLD^12^.

Steatosis has been established to originate in zone 3 in a majority of adult MASLD^1^, supporting the central role of zonal identity in the pathogenesis of this disease. Nonetheless, the specific mechanisms remain elusive. Classically, FAO has been considered to be a zone-1 process, while de novo lipogenesis has been assigned to zone 3^1^. However, more recent transcriptomic and spatial studies have identified PPARα-linked metabolic programs and reported enrichment of fatty acid β-oxidation genes and pathways^23^, and more broadly, enrichment of lipid catabolic pathways in zone 3^23,32,33^, consistent with our findings. A distinction between mitochondrial and peroxisomal β-oxidation has been proposed as one resolution to this paradox^34^, consistent with the necessity of peroxisomal β-oxidation as an input to bile acid synthesis, a zone-3 function. A more in-depth mechanistic study to identify other targets beyond the reported expression of peroxisomal fatty acid oxidation gene Acox1 in zone-3 will be essential in the future. Our findings of increased context-dependent capacity for lipid utilization in HSC-Wls-KO mice are consistent with a lipid metabolism program in zone-3, which is highly adaptive to nutritional state. As a master regulator of the fasting response which is responsible for maintaining lipid, glucose, and protein homeostasis^35–37^, the implication of PPARα in this axis is consistent with the interpretation of zone-3 as a metabolically adaptive program.

β-Catenin and lipid metabolism are closely intertwined with functional consequences. Hepatocyte β-catenin loss has been shown to impair FAO, disrupt PPARα signaling, and increase susceptibility to diet-induced steatohepatitis^38–40^, consistent with both the zone-3 lipid utilization program we identify and the inflammatory phenotype of our HC-Ctnnb1-KO mice. However, transgenic expression of hepatocyte β-catenin resulted in pericentral steatosis under high-fat feeding^38^, in contrast with our Akt-driven model in which acute β-catenin activation limited lipid accumulation. This tension remains unresolved and highlights the nuanced and context-specific metabolic functions of β-catenin which may be further elucidated in future studies. Further, additional targets of Wnt pathway beyond PPARα using shotgun metabolomics and lipidomics integrated with spatial transcriptomics may in future reveal advantages of modulating the master regulator that controls entire zone-3-program over PPARα which control individual processes like FAO. Indeed, several therapies for MASH currently in clinical trials such as pan-PPAR agonists, FGF21 analogues, and combinations of GLP1, GIP and glucagon, all have roles beyond targeting a single process. Wnt agonism is also advantageous in the context of MASH due to its pronounced negative impact on inflammation, a key hallmark of disease pathogenesis and progression. Wnt-β-catenin active cancers including HCCs are immune-cold and β-catenin suppression induces tumor wide immune cell activation through modulation of NF-κB, IRF2 and POU2F1^25^. Conversely, through either directly binding and suppressing NF-κB^41^, or heretofore undetermined mechanisms for IRF2 and POU2F1^42^, β-catenin activation through Wnt agonism, suppresses immune responses which is evident in our study, also in the context of MASH in both preclinical mouse models as well as in LAMPS using human cells. It should also be noted that one of the FDA approved MASH therapeutic resmetirom, which showed benefit in MAESTRO-NASH trial, is a thyroid hormone receptor β analogue^43^. Our previous studies have shown another similar small molecule GC-1 or sobetirome and tri-iodothyronine hormone to activate Wnt-β-catenin pathway directly as well as through stimulating Wnt secretion from liver endothelial cells^44,45^.

Adaptive programs may become pathological when constitutively activated. Our data indicate that the pericentral metabolic program supporting adaptation to nutrient stress is retained, rather than erased, in β-catenin-driven tumors, where it is associated with enrichment for fatty-acid oxidation programs. Notably, tumors lacking coherent zonal identity showed the poorest survival across multiple metrics, suggesting that the organization of zonal identity, and not merely its orientation, carries prognostic information. Whether there is any relationship between HCC etiology and zonation phenotype is not known. *CTNNB1* mutation, which we show is strongly correlated with zone-3 HCC, has not been found to be associated with MASH etiology^11,46^, though future work may be able to decipher whether the zone-3 to zone-1 shift in MASLD may predispose subsequent tumors to a zone-1 phenotype. Recent work has found that zonal identity is an important determinant of tumorigenic potential in HCC^11,47^. Whether the zonal identity of HCC progenitor clones determines eventual zonal subtype, or whether this phenotype is acquired based on hepatocyte-intrinsic evolution or tumor microenvironmental factors, is an open question.

Our investigational pharmacologic studies support that the same zone-3 metabolic program which supports adaptation in MASLD and tumor fitness in HCC may be either activated or inhibited to provide therapeutic benefit, depending on the context. In a genetic model of zone-3 HCC, PPARα antagonism reduced tumor burden, consistent with prior work showing that β-catenin-activated tumors are dependent on fatty-acid oxidation.^22^ Conversely, Wnt agonism in mice with established diet-induced disease reduced pericentral lipid accumulation, though only under dietary conditions permissive for recovery. In a human hepatocyte liver-on-chip system, the same agonist reduced inflammatory cytokine secretion in a model of early metabolic syndrome and improved viability and macrosteatosis under later inflammatory conditions. It should be noted that our findings in the improvement of MASH phenotype are not unequivocal when we force Wnt agonism using FL6.13 in both mice and LAMPS. We attribute this to limited availability of the reagent which requires further dose and frequency optimization. Nonetheless, this is the first proof-of-concept of forcing Wnt signaling and zone-3 as a potential MASH therapeutic. Further, the stage dependence of these responses is itself informative, in that benefit tracked with conditions under which pericentral function could plausibly be restored.

Together, these findings support a view of hepatic zonation as dynamic metabolic organization rather than a fixed anatomical blueprint. Chronic metabolic disease alters not only which metabolic programs the liver expresses, but where coordinated hepatocyte programs are deployed across the lobule, and this arrangement in turn shapes how the liver withstands metabolic stress. Recognizing zonal identity as a dynamic and tunable parameter orchestrating hepatic function lays the groundwork for both understanding the role of zonation remodeling in disease pathogenesis and identifying opportunities for therapeutic intervention.

### Limitations of the study

First, our human analyses are observational and establish an association between disease progression and remodeling of pericentral identity rather than demonstrating that this remodeling initiates disease. Further, we recognize that zone 3 is a complex program with numerous and diverse downstream functions. Manipulation of β-catenin or Wnt secretion therefore raises the likelihood of complex, nonlinear interactions which may demonstrate a high degree of sensitivity to context-dependent factors, including mouse strain and injury model. In particular, the pathway connecting HSC-derived Wnt secretion to remodeling of hepatocyte spatial state warrants further investigation, as does the reconciliation of this phenotype with prior assessments of the same model. We note that our earlier HSC-Wls-KO study, assessed using glutamine synthetase as a single pericentral marker, did not detect a change in zonation, and this deletion did not alter liver injury or fibrosis following CCl_4_ injury or bile duct ligation^8^. More recent work in collaboration with Sugimoto et al. also failed to demonstrate a change in zone 3 in this model^7^, differences which are unexplained but may reflect variation in genetic background, housing conditions, or tissue staining and analysis. In addition, the systemic phenotype of HSC-Wls-KO mice may not be entirely hepatocyte-intrinsic, as an expanded bile acid pool can independently raise energy expenditure^48^. In light of these limitations, HSC-Wls-KO is used in this study as a zone 3 gain-of-function model rather than a definitive conclusion on the role of HSC-derived Wnts, and observations from this model are interpreted cautiously.

In addition, our HCC cohort profiled by spatial transcriptomics is small, and the association between zonated phenotype and survival depends on the application of signatures derived from a small sample of spatial transcriptomics to a larger sample of bulk RNA sequencing data. Finally, the pharmacologic studies are exploratory and lack genetic validation of the therapeutic targets.

## Methods

### RIZLiver Algorithm

RIZLiver begins by assuming that each cell is positioned within the hepatic lobule using a simplified 9-layer coordinate system, from the portal triad to the central vein, whereby gene expression is assumed to be probabilistically influenced by lobule layer. This approach is implemented as a Bayesian model with the following formulation for each gene *g* within cell *j* positioned at layer *k*:

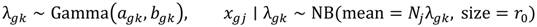

Where *x_gj_* is the observed count of gene *g* in cell *j*, *N_j_* is the library size of cell *j*, *λ_gk_* (expression rate) is the proportion of transcripts of gene g relative to all transcripts in layer *k*, and *r*_0_ is the method of moments global size estimate.

For the Bayesian prior, spatial transcriptomic data from human and mouse liver was manually annotated using spatial visualization of zonation markers (such as *GLUL* and *HAL*) in 5-6 areas of high confidence for each dataset. Three human and two mouse datasets were manually annotated. For each dataset, annotated gene expression data were averaged over a predefined number of discrete regions (varying in granularity based on the platform) and then interpolated into the shared 9-layer coordinate system from the portal triad to the central vein. To pool data across reference sets, the mean and variance for each gene/layer combination across sources were calculated and used as the Gamma prior’s center and width, respectively. (*a_gk_* and *b_gk_* above are calculated from the mean and variance as 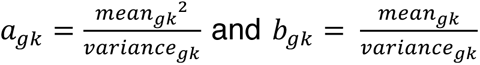). Only data from a single source, the Liver Cell Atlas, was used as a reference for simulated and real-world evaluations (human and mouse). Therefore, the Gamma prior for evaluations was determined by 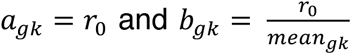. Up to 98 human and 134 mouse landmark reference genes are used for inference depending on availability in the user-provided matrix.

After computing the prior, Bayesian inference is then performed for an initial estimate of lobular position of cells/spots in the user-supplied baseline matrix as ∑*_k_ k P*(*k* ∣ *x*_j_) for each cell *j*. The expression rate *λ_gk_* then undergoes a Bayesian update given the user-supplied baseline data with the initial position estimates, p(λ_g*k*_ ∣ data), effectively calibrating the ground-truth gene zonation profiles to the user-supplied data, which may have batch- and platform-specific differences from the reference data. Finally, the model is refined by recalculating the posterior mean layer ∑*_k_ k P*(*k* ∣ *x*_j_) using the updated expression rate, yielding a layer assignment as a continuous value in the form of a weighted average over posterior probabilities for each layer. These values are then rescaled into a [0, 1] coordinate space via percentile rank, indicating relative position from the portal triad to the central vein.

Trained models using the inference process above can then be applied to additional user-supplied, comparison samples. The trained model is applied to estimate the lobular position based on their count matrices. Their weighted average posterior layers are rescaled into the same [0, 1] coordinate space using the same percentile rank transformation mapped to the baseline distribution via empirical interpolation; comparison sample values which fall outside this range are clamped into the [0, 1] space. As a result, the lobular position estimates in comparison samples are expressed relative to the baseline distribution, rather than their own distributions, allowing RIZLiver to reflect overall changes in the distribution of zonal identity in perturbed samples.

Zonal identity scores are calculated as the log likelihood marginalized over the nine lobule layers, normalized by total landmark gene counts. These scores are normalized to a [0, 1] scale using the percentile rank of baseline scores. Identity score plots, therefore, summarize the cumulative distribution of these cell-or spot-level scores at the sample level, relative to baseline.

By separately annotating both human and mouse datasets, RIZLiver provides species-specific inference of zonal identity, with species specified by the user at run time. The full source code and usage instructions are available at https://github.com/tyleryasaka/RIZLiver.

### Zonation Inference Evaluations

Simulated gene expression matrices were constructed from human and mouse spatial transcriptomics data from the Liver Cell Atlas^49^, which were manually annotated to indicate the ground-truth zonation axis. Average gene expression values across discretized zonated regions defined by these manual annotations were interpolated into a 100-position ground-truth vector. Zonated genes were identified as genes with statistically significant (adjusted p-value < 0.01) Spearman correlations with the manually annotated zonation axis while all other gene expression values were randomly shuffled at baseline to control for non-zonation artifacts in the dataset.

Usage of the same dataset for RIZLiver reference data and simulation ground truth was deliberate in order to perform a tightly controlled experiment. Therefore, baseline simulations evaluate the scenario where the ground truth is perfectly captured by the reference data, while adding axis rotation allows for varying discrepancy between the ground truth and reference data to be introduced in a controlled manner. To ensure fairness, algorithms which use landmark genes (Halpern, Pseudogradient, and AddModuleScore) used the top three periportal and top three pericentral zonated genes, as determined by Spearman correlation with the manually annotated axis in the ground-truth dataset, as landmark genes during evaluation.

Implementation of alternative methods for benchmarking was performed as described in the respective papers^13,14,50^. The more generic pseudotime and ModuleScore methods represent multiple papers^23,51–54^. For ModuleScore, the AddModuleScore method from the Seurat library was called using landmark genes as the feature set to assign a score to each cell. For pseudotime, a one-dimensional trajectory was computed from the PCA embeddings of the Seurat object as previously described^55^, with landmark genes used for directional orientation of the trajectory. For all methods, the final position estimates were transformed to the same [1, 100] scale as the ground truth using percentile ranking so that relative, rather than absolute, estimates were evaluated.

Technical variability was modeled as sequencing depth where each cell’s counts were scaled by a factor drawn from a log-normal distribution centered around the specified sequencing depth, while biological variability was modeled as overdispersion of the expression rates for each gene/cell combination following a negative binomial distribution, with a Poisson distribution used when overdispersion = 0.

Axis rotation was introduced by first calculating a signal vector representing each gene’s loading on the zonation axis, then calculating a random orthogonal decoy vector of equal magnitude. A Givens rotation was performed between the signal vector and the decoy vector at the specified axis rotation. The latent factor was implemented as a randomly generated independent axis where each cell was assigned a score drawn from a normal distribution, and each gene was given a coefficient randomly drawn from a normal distribution with standard deviation determined by the latent factor strength. The product of the axis score and the gene coefficient was added to the log expression for each gene/cell combination.

Simulation experiments were performed separately for human and mouse with 10 trials for each parameter combination. In each trial, 1000 simulated cells were generated and distributed over the 100 simulated positions across the lobule from the portal triad to the central vein (10 cells per position). The gene expression of each cell was determined from the ground-truth matrix, with varying expression perturbations based on the experimental parameters as described above. Positions of simulated cells estimated from the evaluated inference algorithms, including RIZLiver and comparators, were compared with the ground-truth positions to determine the Spearman correlations and root mean squared error (RMSE).

For real-world dataset evaluations, ground-truth positions were determined based on relative locations within manually annotated segments, using spatial visualization of zonation markers, spanning the portal triad to the central vein with benchmarking metrics measured as above.

### Mice

All animal experiments were conducted in accordance with the guidelines of the National Institutes of Health and were approved by the Institutional Animal Care and Use Committee at the University of Pittsburgh. Mice were maintained under standard housing conditions with a 12-hour light–dark cycle and provided ad libitum access to regular chow, without environmental enrichment. Hepatic stellate cell (HSC)-specific Wntless knockout (HSC-Wls-KO) mice were generated by crossing C57BL/6 *Lrat*-Cre mice (kindly provided by Dr. Robert Schwabe^56^) with homozygous floxed *Wls* mice (*Wls*^loxp/loxp^). *Lrat*-Cre^+/-^; *Wls*^loxp/WT^ offspring were subsequently crossed with *Wls*^loxp/loxp^ mice to obtain *Wls*^loxp/loxp^; *Lrat*-Cre^+/-^ mice (HSC-Wls-KO) on a C57BL/6 background. Littermate controls (Con) consisted of *Wls*^loxp/loxp^; *Lrat*-Cre^-/-^ or *Wls*^loxp/WT^; *Lrat*-Cre^-/-^ mice. Efficiency and specificity of Cre-mediated recombination in HSCs were validated by crossing *Wls*^loxp/loxp^; *Lrat*-Cre^+/-^ mice with ROSA26^LSL-EYFP^ reporter mice. Genotyping was performed using PCR with primer sequences and expected product sizes provided in **Tables S2 and S3**.

To generate hepatocyte-specific β-catenin knockout (HC-Ctnnb1-KO) or control mice, 8-10 weeks old *Ctnnb1^flox/flox^* mice were injected intraperitoneally with 1×10^12^ genome copies of AAV8-TBG-Cre or AAV8-TBG-GFP (Cre recombinase or GFP driven by the hepatocyte-specific thyroxine-binding globulin promoter), followed by 12-day washout period.

### Single-nucleus RNA sequencing (snRNA-seq)

For snRNA-seq, fresh frozen liver tissue was collected and sent to Novogene for analysis. Both the wild-type and the knock-out libraries were measured by snRNA-seq using one mouse each. Software Cell Ranger^57^ count function was applied to each library to align the sequence file to mouse reference genome mm10 and to quantify the gene count per nucleus. After pre-processing, the R package Seurat^58^ was applied to the gene count data for downstream analysis.

We first performed filtering step and only keep the nuclei with higher than 500 genes expressed, lower than 6000 genes expressed, less than 12000 reads and less than 10% of mitochondria gene expression. Next, we applied the *SCTransform* function to normalize each library and integrate both libraries by top 3000 high variable features. After these, dimensionality reduction was performed and single nuclei were visualized by Uniform Manifold Approximation and Projection (UMAP) algorithm. Finally, nuclei were clustered based on their gene expression pattern, and known marker genes were applied to annotate the cell population with cell types found in the liver.

### Single-nucleus RNA sequencing using 10x Genomics Chromium Flex

Flash-frozen liver tissues were processed by the Single Cell Core at the Pitt Center for Advanced Genomics for nuclei isolation and processing on the 10x Genomics Chromium Flex platform per the manufacturer’s protocol. These services are available through the Pittsburgh Liver Research Center (1P30DK120531). QC was performed by filtering out cells with fewer than 500 total features. Seurat clusters were annotated based on cell type. RIZLiver was applied to hepatocytes to infer zonal identities, trained on control hepatocytes and applied to HC-Ctnnb1-KO.

### Resolve Molecular Cartography^TM^ Spatial Transciptomics

Resolve Molecular Cartography^TM^ was performed as previously described^6,59^. Briefly, formalin-fixed paraffin-embedded (FFPE) tissue sections from one HSC-Wls-KO and one litter matched control mouse were sent for Molecular Cartography^TM^. QuPath was used to perform cell segmentation based on nuclear DAPI staining of the slides, followed by single-cell spatial transcriptomic analysis using Seurat. QC was performed by filtering out cells with fewer than 10 total transcripts. Seurat clusters were annotated based on cell type. RIZLiver was applied to hepatocytes to infer zonal identities, trained on control hepatocytes and applied to HSC-Wls-KO.

### Xenium In Situ Spatial Transcriptomics

Xenium In Situ spatial transcriptomics was performed by the Translational Oncologic Pathology Services (TOPS) core at the University of Pittsburgh Medical Center (UPMC) using FFPE tissue sections from resected human HCC and one healthy control liver. These services are available through the Pittsburgh Liver Research Center’s Cellts, Tissues and Models Core as well as Omics and Systems Biology Core. Samples were mounted onto two slides which were imaged on a single Xenium run. QC was performed by filtering out cells with fewer than 10 total transcripts and/or fewer than 5 total features. Data was processed using Seurat v5 integration with Harmony batch correction for Xenium slide.

### Indirect Calorimetry and Body Composition Measurements

The Promethion Multiplexed Metabolic Cage System (Sable Systems) by the University of Pittsburgh Center for Metabolism and Mitochondrial Medicine was used to perform indirect calorimetry as previously described^60^. Male Con (n=7) and HSC-Wls-KO (n=7) mice were individually placed in cages and allowed to acclimate for 24h. To establish a baseline metabolic phenotype with high statistical power, metabolic cage studies were limited to male mice to minimize biological variability and facilitate robust detection of metabolic effects. Mice were then subjected to 48h at ambient room temperature with 12h light/12h dark cycles for analysis and had access to food and water. Total activity data from one HSC-Wls-KO mouse were excluded due to an unresolvable infrared sensor artifact. Body composition including fat and lean mass was determined using EchoMRI.

### Mitochondrial respirometry

Mitochondrial enriched supernatants were prepared using fresh liver samples from 3 Con and 3 HSC-Wls-KO male mice, as previously described^60^. Respirometry was then performed using Oroboros O_2_K High-Resolution Respirometer on these mitochondrial enriched samples. Respiration was measured in MiR05 buffer at 37°C under constant mixing in a sealed, 2-ml chamber. Sequential additions of substrates and inhibitors were as follows: pyruvate/malate (5 and 2 mM) or palmitoylcarnitine (40 µM), adenosine diphosphate (ADP, 0.2 mM and 2.0 mM); and glutamate (10 mM); succinate (10 mM) carbonyl cyanide 4-(trifluoromethoxy)phenylhydrazone (FCCP; titrations of 0.5 µM until maximal respiration reached), and rotenone (0.5 µM). Respirometry data were normalized to total protein measured by BCA protein assay.

### Western Diet

Littermate- and gender-controlled Con and HSC-Wls-KO mice, or control and HC-Ctnnb1-KO were fed a western diet for 3 months according to the protocol described in Rai et al^61^. Body weights were measured once per month, and mice livers and serum were harvested at the end of the third month.

### Bile acid mass spectrometry

Bile acid analysis was performed as previously described^62^. Briefly, liver tissues from WT and HSC-Wls-KO mice (n = 3) were homogenized in water (100 mg tissue per 500 µl water). Methanol–acetonitrile (1:1, v/v; 300 µL) was added to 100 µL of each liver homogenate. Samples were then vortexed for 2 minutes and centrifuged at 15,000 rpm for 10 minutes. An aliquot (2 µL) of the supernatant was injected into the ultra-performance liquid chromatography (UPLC) coupled with a SYNAPT G2-S quadrupole time of flight mass spectrometry (QTOFMS) (Waters Corporation, Milford MA) for analysis. The Acquity UPLC BEH C18 column (2.1 x 100 mm, 1.7 µm) was used for metabolite separation. The mobile phase gradient details are described by Jiang et al^63^. The QTOFMS system was operated in a negative high-resolution mode with electrospray ionization^63^. Bile acid species were then quantified by measuring their relative abundance as the area under the curve for each species based on comparisons with standards^64^.

### Immunohistochemistry

Livers were fixed in 10% buffered formalin for 48 hours and paraffin embedded. Blocks were then cut into 4µm sections. Sections were deparaffinized and stained as described in previous studies^6^. Primary antibody dilutions, incubation times, and biotinylated secondary antibodies are described in **Table S4**. Images were taken on the Zeiss Axioskop 40 inverted brightfield microscope.

### Real Time PCR

RNA was extracted from whole liver tissue using TRIzol (Thermo Scientific, 15,596,026). Nucleic acid was separated using phenol-chloroform extraction. RNA was reverse transcribed to cDNA with SuperScript III (Invitrogen, 18080-044). Real-time PCR was performed in technical duplicates on a StepOnePlus Real-Time PCR System (Applied Biosystems, 4-376-600) using Power SYBR Green PCR Master Mix (Applied Biosystems, 4-367-660). The housekeeping gene, Rn18s, was used to normalize target gene expression. Fold change was calculated with the ΔΔ-Ct method. The list of primers used is provided in **Table S5**.

### LAMPS assembly and maintenance

Hepatocyte-only Liver Acinus Microphysiological Systems (LAMPS) were assembled and maintained as previously described using the Fluidic 557 Reaction Chamber Chip (Microfluidic ChipShop #557)^65^. Briefly, LAMPS were built using primary cryopreserved human hepatocytes (2.70–2.85 × 10⁶ cells/mL) (ThermoFisher lot hu8450). Prior to cell seeding, the device interiors were coated with 100 μg/mL bovine fibronectin (Sigma-Aldrich, F1141) and 150 μg/mL rat tail collagen I (Corning, 354249) in PBS. To ensure full coverage of fluidic pathways, chambers, 90–100 μL of media or cell suspension was injected per device. After hepatocyte attachment ∼ 6hours cells were overlaid with decellularized porcine liver extracellular matrix (LECM)^65,66^ made up in Early Metabolic Syndrome media. On the following day, the devices were then overlaid with 1.5 mg/mL rat-tail collagen I (Corning) and maintained under perfusion with different media conditions for 4 days at a flow rate of 5 μL/h, mimicking zone-3 oxygen tension as previously described^67^.

### Formulation Early Metabolic Syndrome and Late Metabolic Syndrome Media Formulation and FL6.13 intervention

MASLD/MASH disease phenotypes in LAMPS are driven using our previously published media formulations^68,69^. These media formulations were developed using glucose-free Williams E base medium (ThermoFisher, ME18082L1) supplemented with varying levels of glucose (MilliporeSigma, G8644), insulin (ThermoFisher, 12585014), and glucagon (MilliporeSigma and molecular drivers of fibrosis including TGF-β1 (ThermoFisher, PHG9214) and lipopolysaccharide (MilliporeSigma, L2654). FL6.13 was administered at a concentration of 50 nM in LAMPS.

### Confocal imaging and analysis

Imaging was conducted on the Phenix High-Content Imaging system (Revvity) using a 20×/0.4 NA air objective. Z-stacks covering 35 μm depth (5 μm steps) were acquired across 7 × 15 adjacent fields (total area: 44 mm²). All images were collected using consistent exposure and laser power settings to maintain signal intensity within 50–90% of the dynamic range. Image analysis was performed using Harmony software (Revvity, v5.1).

### CK-18 Immunofluorescence and LipidTOX staining and Quantification

Cells were fixed with 4% paraformaldehyde (ThermoFisher Scientific, AA433689M) in PBS for 30 min then washed twice with PBS for 10 min at room temperature. Following fixation, LipidTOX Deep Red Neutral Lipid Stain (1:500, Invitrogen, H34477) and rabbit polyclonal anti-cytokeratin18 (CK-18) antibody (ThermoFisher PA5-14263; 1:200) in PBS were perfused into devices and incubated overnight at 4°C. The following day, devices were washed twice with PBS and then incubated for 2 h with Alexa Fluor Goat anti-rabbit 488 (1:250, Invitrogen, A-11029) secondary antibody and Hoechst (5 μg/mL, Invitrogen, H1399) at room temperature. Devices were washed three times with PBS before imaging. Steatosis was quantified from LipidTOX-stained images using automated segmentation and object detection workflows (Harmony). Nuclei were identified from the DAPI channel, and lipid-positive regions were segmented based on LipidTOX fluorescence (ex: 640 / em: 650-760). Individual lipid droplets between 5 – 30 μm in diameter were detected and counted. Lipid accumulation was reported for both the mean lipid-positive area and the % Macrovesicular steatosis per chamber. The % Macorvesicular steatosis was calculated by determining if there were any lipid vesicles in the cell greater than 8 microns in diameter, then classifying the cell as containing a macrovesicular-containing cell. Then, the percentage of macrovesicular steatosis was determined by the following calculation: the number of macrovesicular-containing cells divided by the total number of cells (non-macrovesicular-containing cells and macrovesicular-containing cells) and multiplied by 100.

### Calcein AM staining and Quantification

For endpoint live cell imaging, cultures were stained with Calcein AM (ThermoFisher C1430) and Hoechst nuclear dye (ThermoFisher 62249). A 1 mg/mL Calcein AM stock solution was prepared in DMSO and diluted 1:1000 in normal fasting medium to yield a final concentration of approximately 1 µM, together with Hoechst (1:2000; ∼9 µM final). Following chip disconnection, 50 µL of staining solution was applied to both the inlet and outlet, and chips were incubated for 45 min at 37 °C. Imaging was performed at 10× magnification using the FITC (Calcein AM) and DAPI (Hoechst) channels. Live cell quantification was performed using an automated image analysis workflow implemented in Harmony (PerkinElmer). The analysis utilized the DAPI (nuclear) (ex: 405/ em: 435-480) and calcein (live cell) (ex: 488 / em: 500-550) fluorescence channels to identify total and live cell populations. Nuclei were segmented using the “Find Nuclei” block set to method M in the Harmony 5.1 software, and calcein-positive regions were detected using optimized intensity thresholding within the “Find Image Region” block. Then nuclei not overlapping with calcein-positive regions were excluded using the “Select Population” block. The percentage of live cell area was determined as:

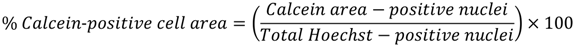

### Multiplex immunoassays

The levels of IL-6, IL-8, and CCL-2, IL-1β, IL-10 and TNFα were determined in efflux collected on day 8 using a custom version of the Human XL Cytokine Performance Panel (R&D systems). Assays were completed according to the manufacturer’s instructions at The University of Pittsburgh Cancer Proteomics Facility Luminex® Core Laboratory on the xMAP platform.

### FL6.13 administration in Western diet-fed mice

Male C57BL/6 mice were fed a Western diet for 12 weeks prior to treatment initiation. The Wnt agonist antibody FL6.13 was administered by intraperitoneal injection at 10 mg/kg every other day, as previously described^6^. Control mice received either control IgG or saline. Mice were divided into two experimental groups. In Group A, mice were maintained on Western diet throughout the treatment period and received a total of five injections (FL6.13, *n* = 4; control, *n* = 3). In Group B, mice received four injections while on Western diet, followed by a switch to normal chow diet and two additional injections prior to tissue collection (FL6.13, *n* = 4; control, *n* = 3). Body weight was monitored during treatment. Liver tissue and visceral adipose tissue were harvested for further analysis.

### Akt-induced hepatic steatosis model and PPARα antagonist treatment

Akt-driven hepatic steatosis was induced by hydrodynamic tail vein injection (HTVI) of plasmids encoding constitutively active Akt alone or in combination with constitutively active β-catenin (S45Y). Mice were sacrificed 14 days after injection for tissue collection. For PPARα inhibition, the PPARα antagonist GW6471 (HY-15372, MedChemExpress) was administered by intraperitoneal injection at 20 mg/kg. Treatment was initiated 24 hours after HTVI and continued every other day for a total of seven injections. Control mice received vehicle injections on the same schedule. At day 14, mice were euthanized and livers were harvested for subsequent analyses.

### Oil Red O staining

Frozen liver sections (10 μm thickness) were prepared using a cryostat and subjected to Oil Red O staining. Lipid accumulation was visualized using Oil Red O solution (ab150678, Abcam) according to the manufacturer’s instructions. Sections were counterstained with hematoxylin, mounted, and imaged using a Zeiss Axioskop 40 inverted brightfield microscope. Lipid droplets were quantified by counting the number of droplets per field of view at 20× magnification.

### Percent area quantification

Four to five random 10x fields per liver sections were captured per animal, excluding artifacts, and analyzed in ImageJ. Percent positive area was calculated as (positive-stained area/total tissue area) x 100 per animal using a fixed threshold applied uniformly across all sections.

### Statistical analysis

Statistical analyses were performed using GraphPad Prism 10.5 (GraphPad Software, San Diego, CA, USA). Data are presented as mean ± SD. All datasets were assessed for normality and homogeneity of variance prior to analysis. Group differences were evaluated using the unpaired Student’s T-test or one-way analysis of variance (ANOVA), followed by Tukey’s multiple-comparisons test where appropriate, unless otherwise specified. Statistical significance was defined as p < 0.05.

## Materials availability

Mouse lines and all unique reagents used in this study are available from the lead contact with a completed Material Transfer Agreement.

## Data and code availability

The snRNA-seq and spatial transcriptomics datasets generated in this study has been deposited in the Gene Expression Omnibus (GEO) under accession number GSE352587. The software and algorithms used for data analysis are described and referenced throughout the STAR Methods section. Additional information required to reanalyze the data reported in this paper is available from the lead contact upon reasonable request.

## Acknowledgements

This work was supported by NIH grants 1R01DK62277, 1R01DK103775, 1R01CA251155, 1R01CA250227 and Senior Vice Chancellor Endowed Chair in Pathobiology and Therapeutics to S.P.M. This work was also funded in part by R01DK124510 to N.K.H.K., R01DK114012 and R01DK119627 to M.J.J., NIH/NIGMS R35GM159862 to S.L., F30CA298277 to T.M.Y., T32EB001026 to T.M.Y. and B.M.L., T32EB001026 and the Health Innovation in Translational Science (HITS) Pilot Award through the University of Pittsburgh Clinical and Translational Science Institute (UL1TR001857) to T.M.Y., B.M.L., Y.-C.C. and S.P.M., F30CA284540 to B.M.L., and R01DK124351 to R.R. This work was also supported in part by the University of Pittsburgh Center for Research Computing and Data through the resources provided (supported by NIH grant S10OD028483) and by NIH grant 1P30DK120531 to Pittsburgh Liver Research Center (PLRC) for services provided by the Cells, Tissues and Models Core and the Omics and Systems Biology Core. This work was supported by a grant-in-aid from the Research Program on Hepatitis from the Japan Agency for Medical Research and Development (AMED JP25fk0210172). The cytokine data was obtained using the UPMC Hillman Cancer Center Biomarkers Facility: Luminex Core Laboratory.

## Author Contributions

N.K.H.K. and S.P.M. conceived the study. H.I., T.M.Y., C.K.K., N.K.H.K., and S.P.M. designed the experiments. H.I., T.M.Y., C.K.K., A.S.-V., S.H., B.M.L., C.K.K., L.M., M.P., A.J., V.M., X.M., A.M.V., R.P.R., S.S., I.S., F.M.B., J.A.B. and A.B. performed the experiments. T.M.Y., H.I., A.S.-V., J.-J.L., J.A.B., M.T.M., J.L., Y.R., S.U., M.J.J., N.K.H.K. and S.L. analyzed the data. T.M.Y., H.I., C.K.K., N.K.H.K., M.T.M, and S.P.M. wrote the manuscript. T.M.Y., B.M.L., N.K.H.K., and S.P.M. acquired funding. R.P.R., I.S., F.M.B., R.R., J.A.B., M.T.M., Y.T., Y.-C.C., M.J.J., S.A., N.K.H.K. and S.P.M. provided resources.

