## Supplemental Figures for "Dynamic zonal identity is a functional determinant of liver disease"

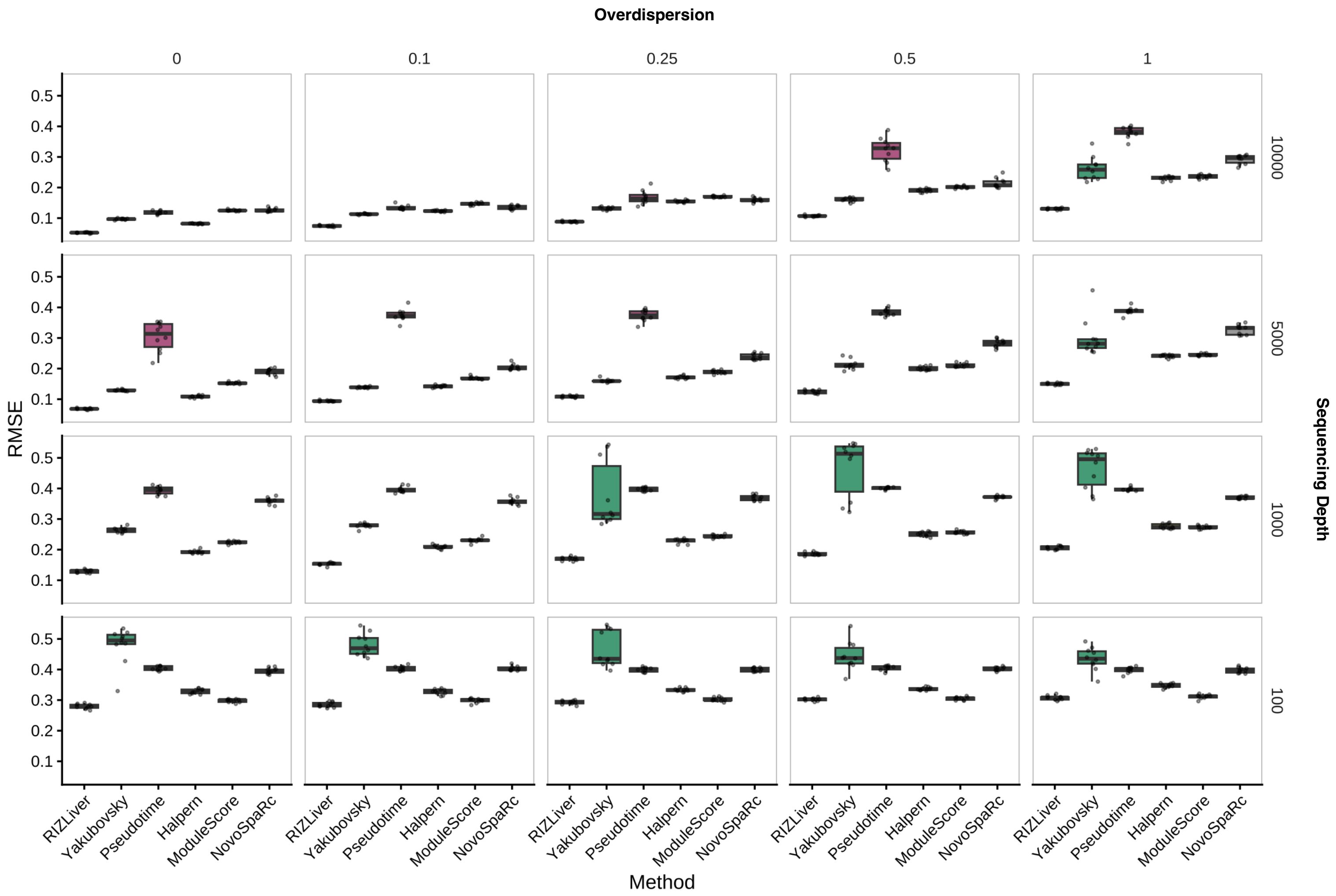


**Figure S1.** Evaluation of zonation inference methods in simulated human experiments across multiple biological variability (overdispersion) and technical variability (sequencing depth) values. Displayed values are average root mean squared error (RMSE) values across 10 simulations.

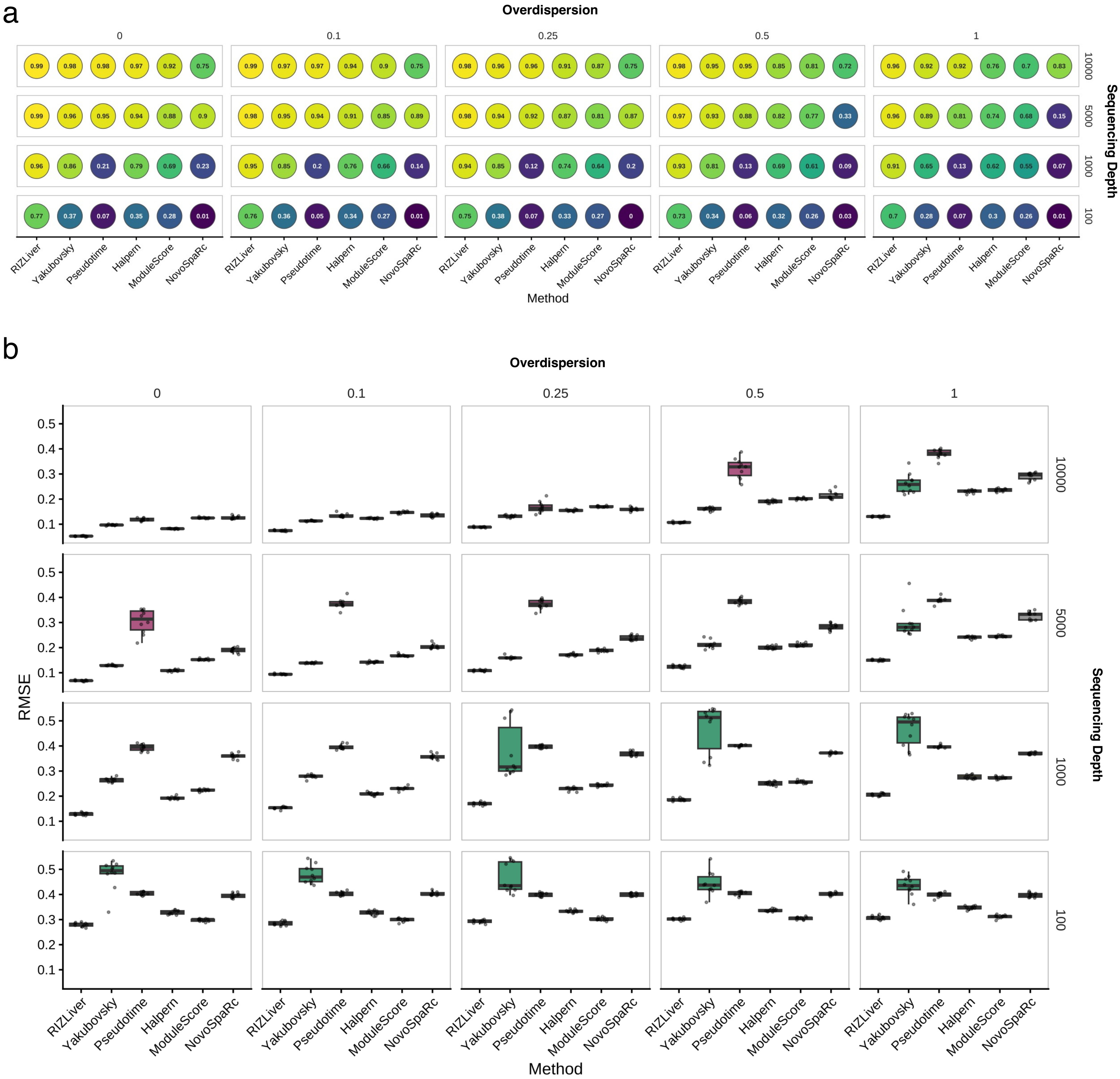


**Figure S2. (a,b)** Evaluation of zonation inference methods in simulated mouse experiments across multiple biological variability (overdispersion) and technical variability (sequencing depth) values. Displayed values are **(a)** average spearman correlation coefficients and **(b)** average root mean squared error (RMSE) values across 10 simulations.

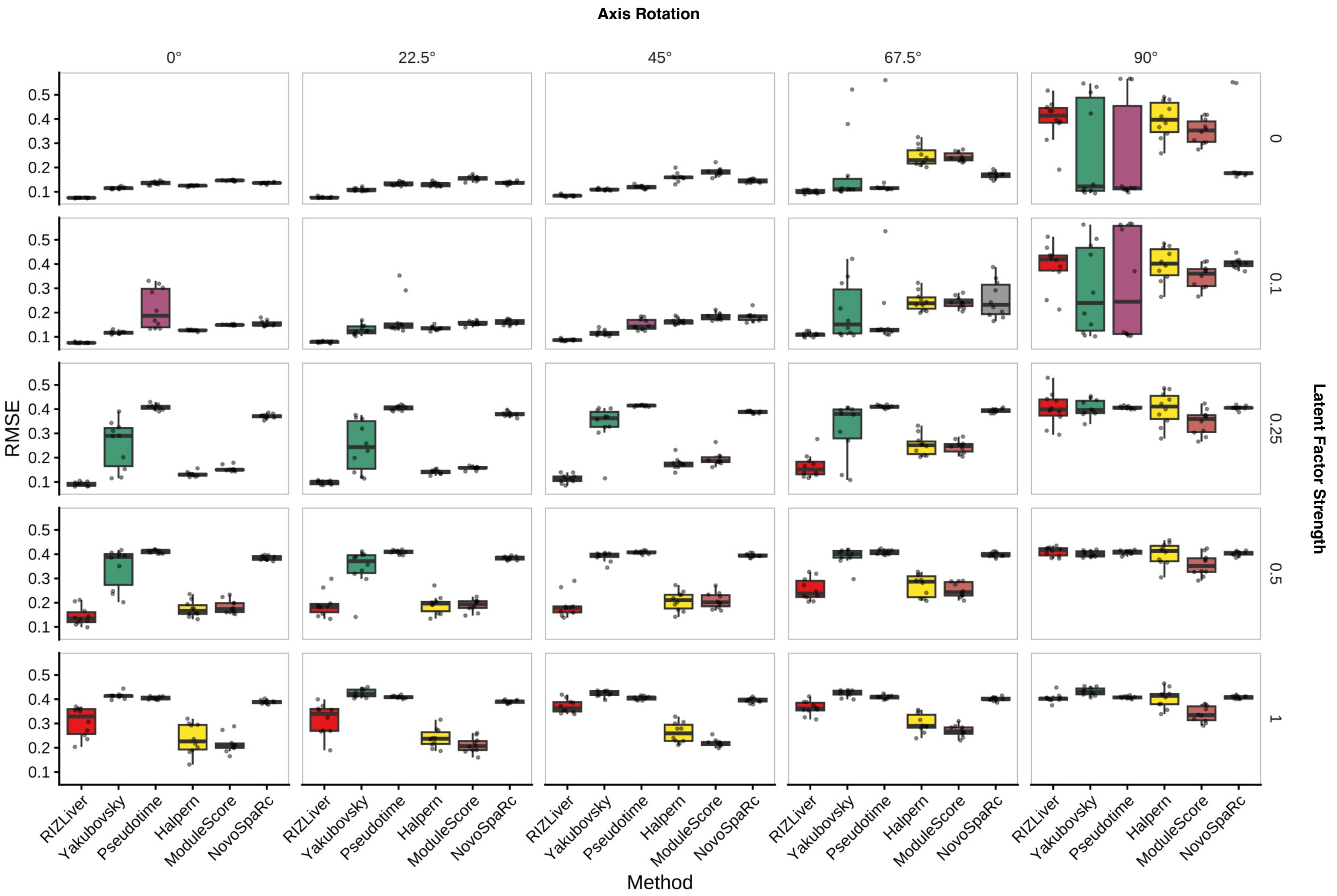


**Figure S3.** Evaluation of zonation inference methods in simulated human experiments across multiple axis rotation degrees and latent factor strengths. Displayed values are average root mean squared error (RMSE) values across 10 simulations.

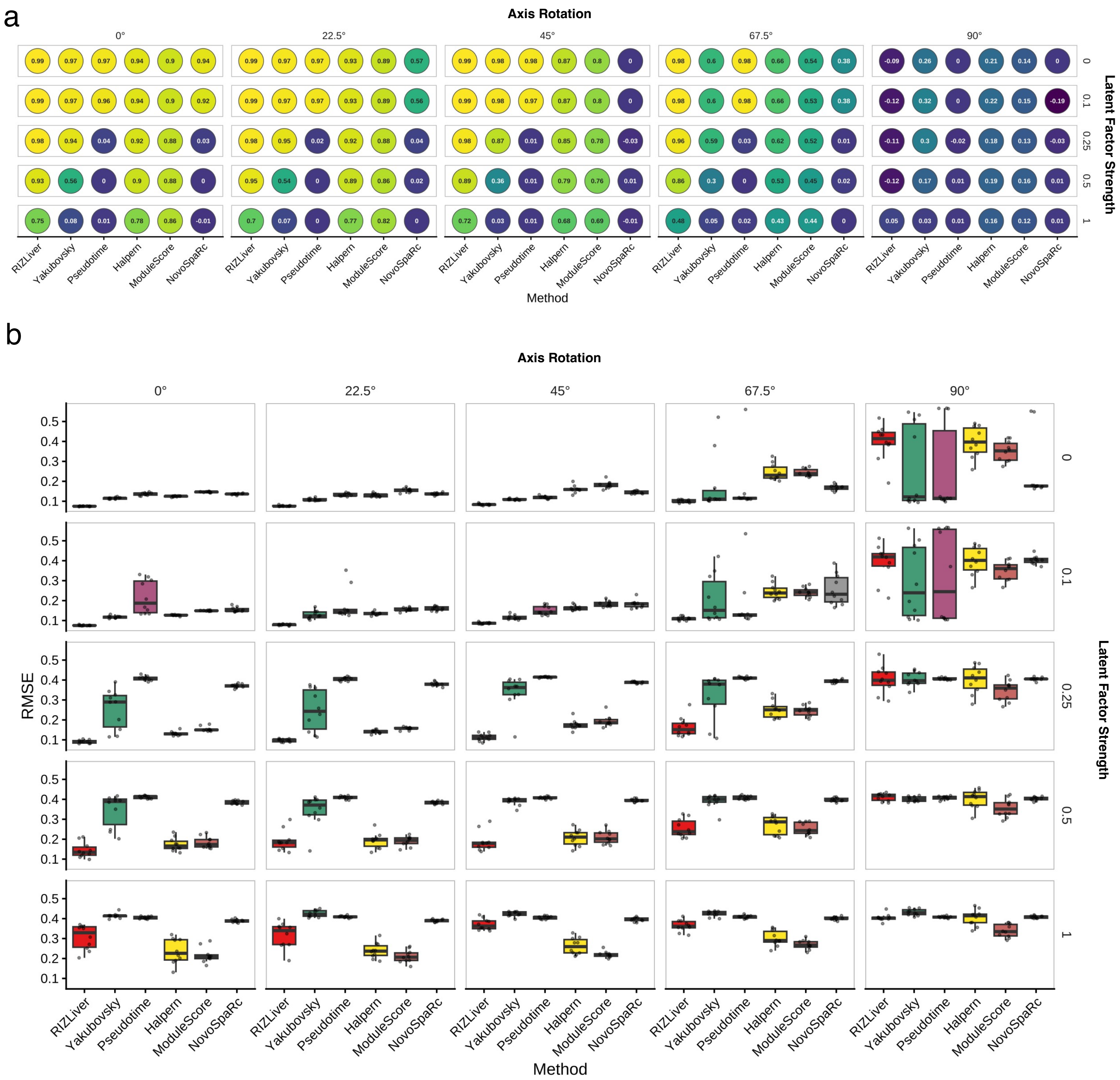


**Figure S4. (a,b)** Evaluation of zonation inference methods in simulated mouse experiments across multiple axis rotation degrees and latent factor strengths. Displayed values are **(a)** average spearman correlation coefficients and **(b)** average root mean squared error (RMSE) values across 10 simulations.

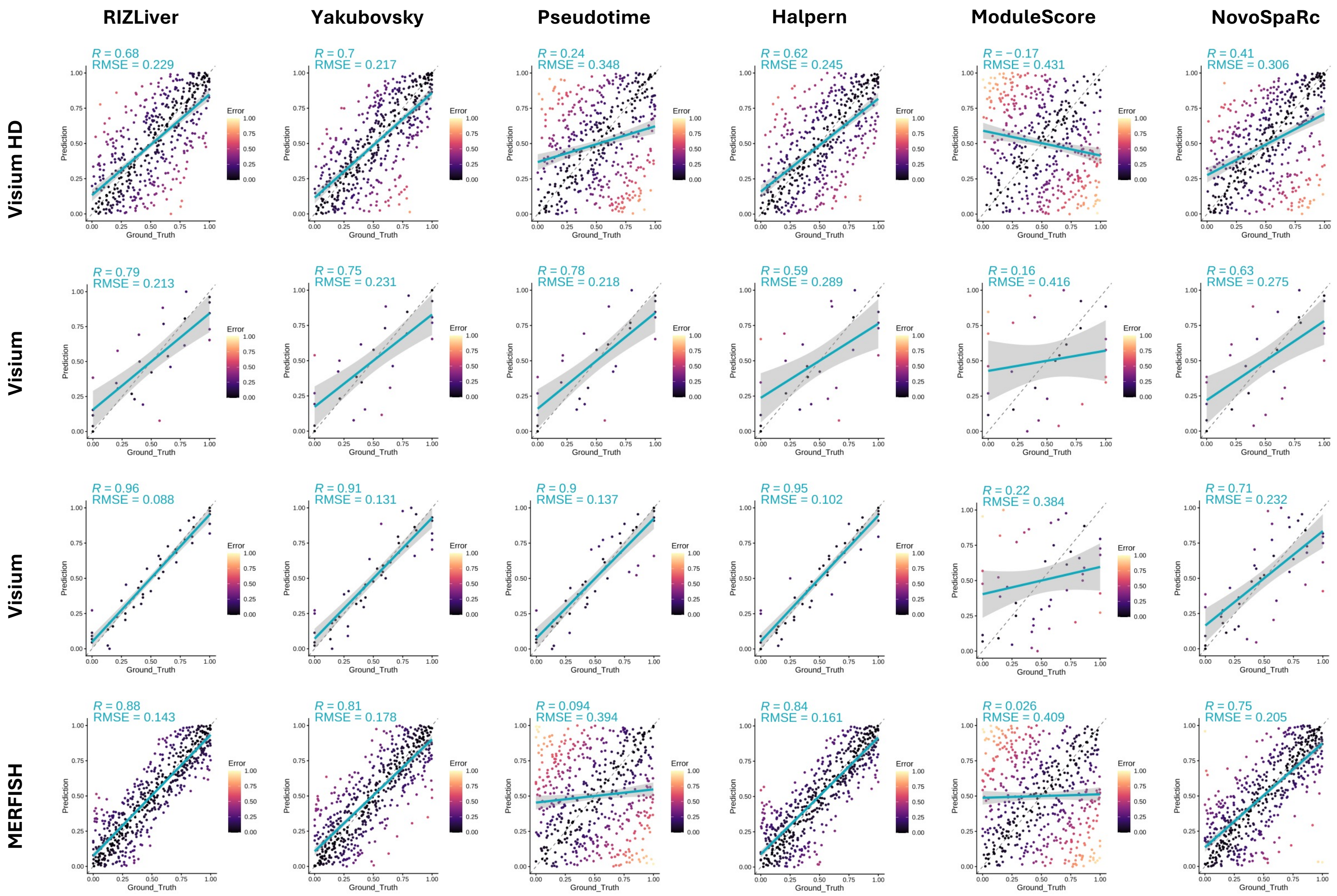


**Figure S5.** Evaluation of zonation inference methods using manually-annotated human spatial transcriptomic datasets. Inferred positions (y-axis) are compared with ground-truth relative locations within the lobule (x-axis) defined by manual annotation. Spearman correlation coefficients and average root mean squared error (RMSE) values are displayed.

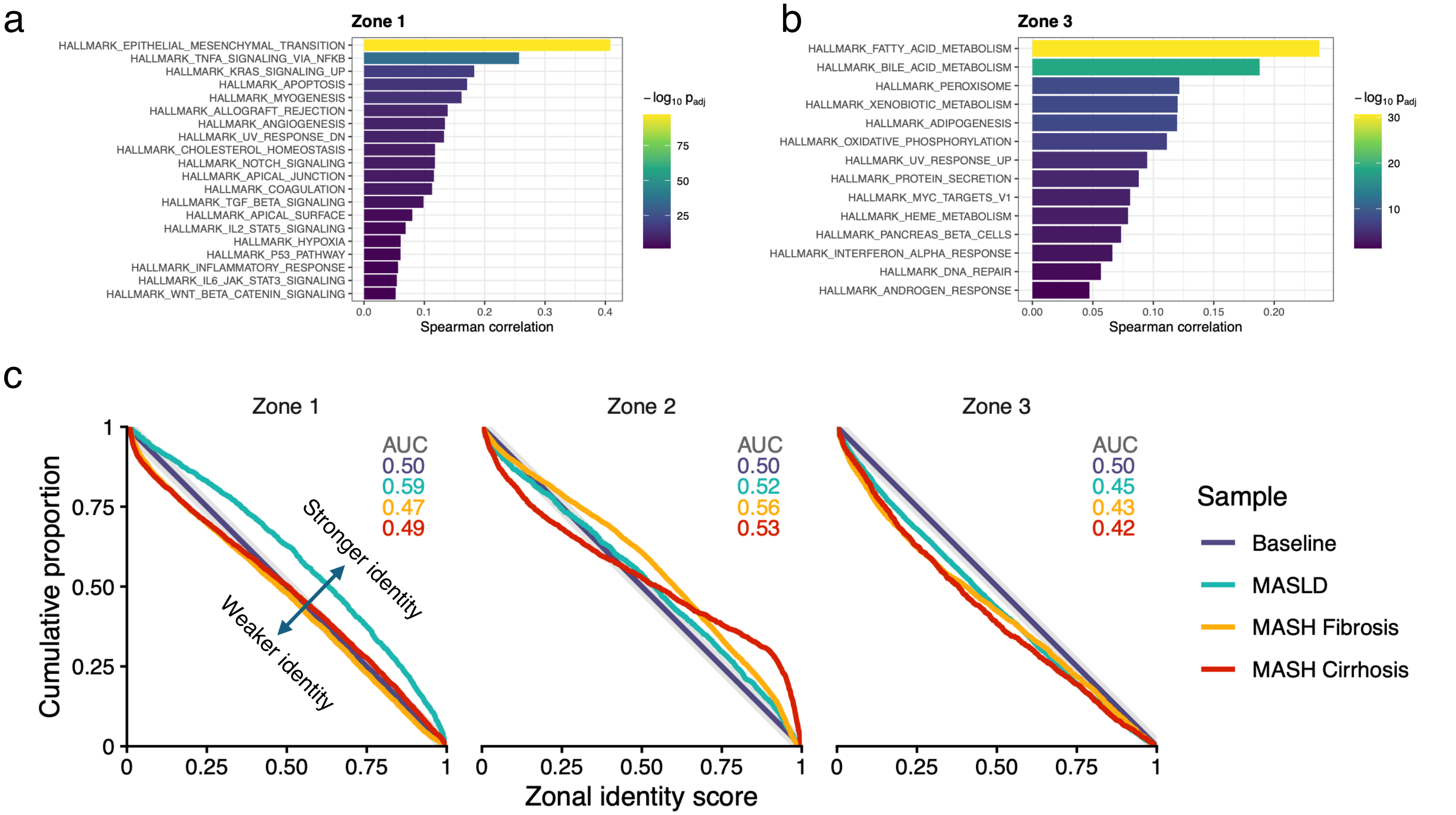


**Figure S6. (a,b)** Top zonated Hallmark pathways as ranked by spearman correlation between inferred zonation and single sample gene set enrichment analysis (ss-GSEA) in Visium spatial transcriptomics of healthy human liver. **(a)** Top zone-1-enriched pathways. **(b)** Top zone-3-enriched pathways. **(c)** Zonal identity scores of cells inferred to be in zones 1, 2 and 3 based on relative lobular position. An AUC above 0.5 indicates overall stronger zonal identity relative to baseline, while an AUC below 0.5 indicates weaker zonal identity relative to baseline.

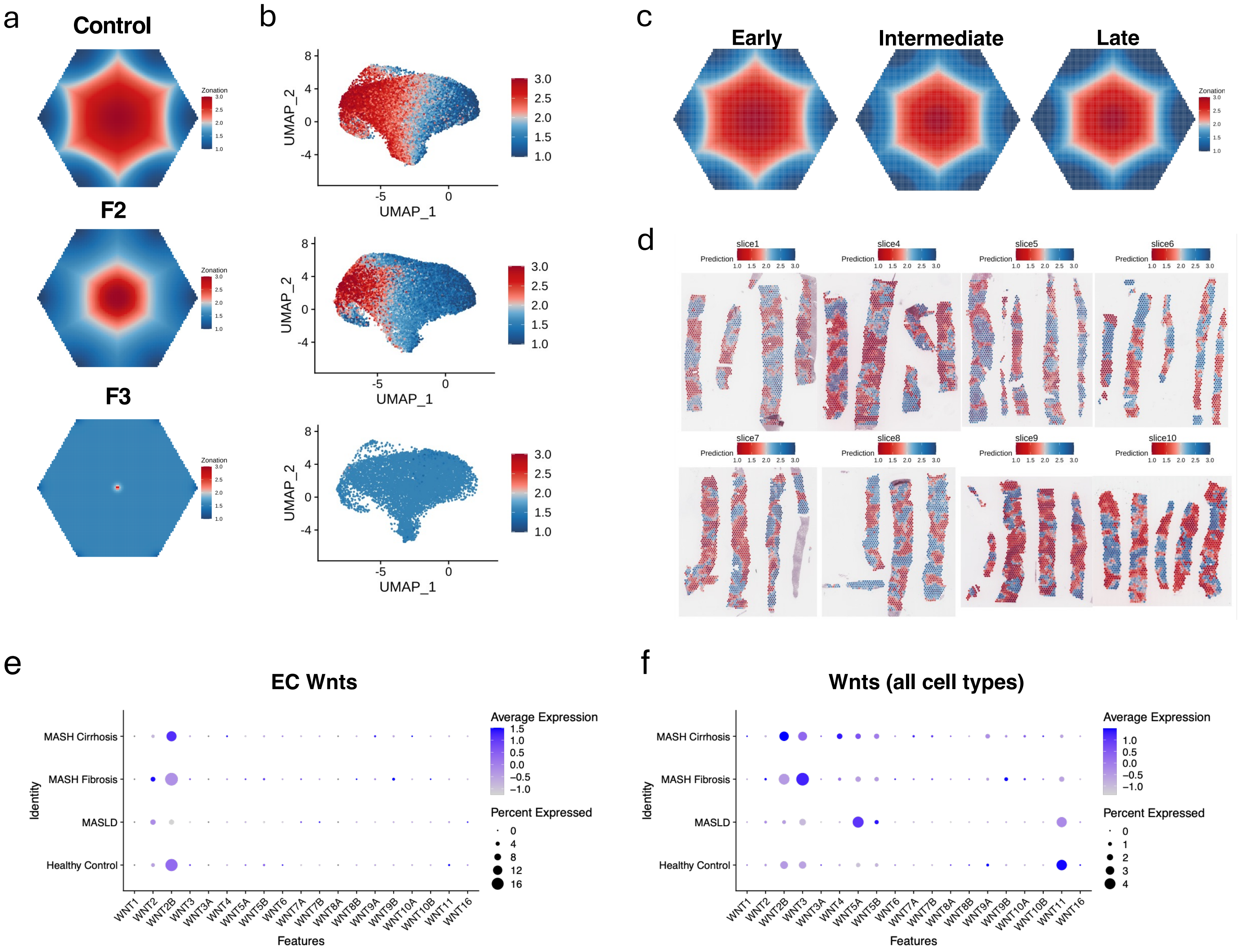


**Figure S7**. **(a, b)** Virtual lobule and UMAP plots of zonal identities inferred using RIZLiver comparing F2 and F3 MASH to control liver in snRNA-seq data. **(c, d)** Virtual lobule and spatial plots of zonal identities inferred using RIZLiver comparing intermediate and late MASLD to early MASLD in Visium spatial transcriptomic data. **(e)** Total expression of Wnt genes in endothelial cell snRNA-seq across stages of MASLD. **(f)** Total expression of Wnt genes in snRNA-seq across stages of MASLD.

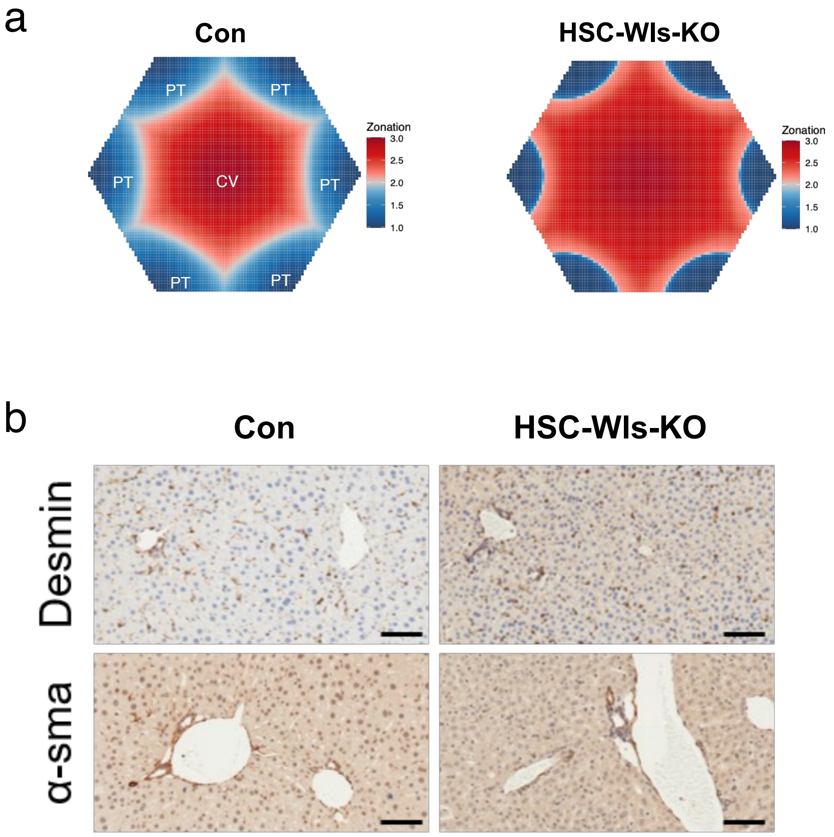


**Figure S8**. **(a)** Virtual lobule plots of zonal identities inferred using RIZLiver analysis of snRNA-seq comparing HSC-Wls-KO versus control. **(b)** Representative immunohistochemical staining of HSC activation markers (Desmin, α-SMA) in HSC-Wls-KO and control livers.

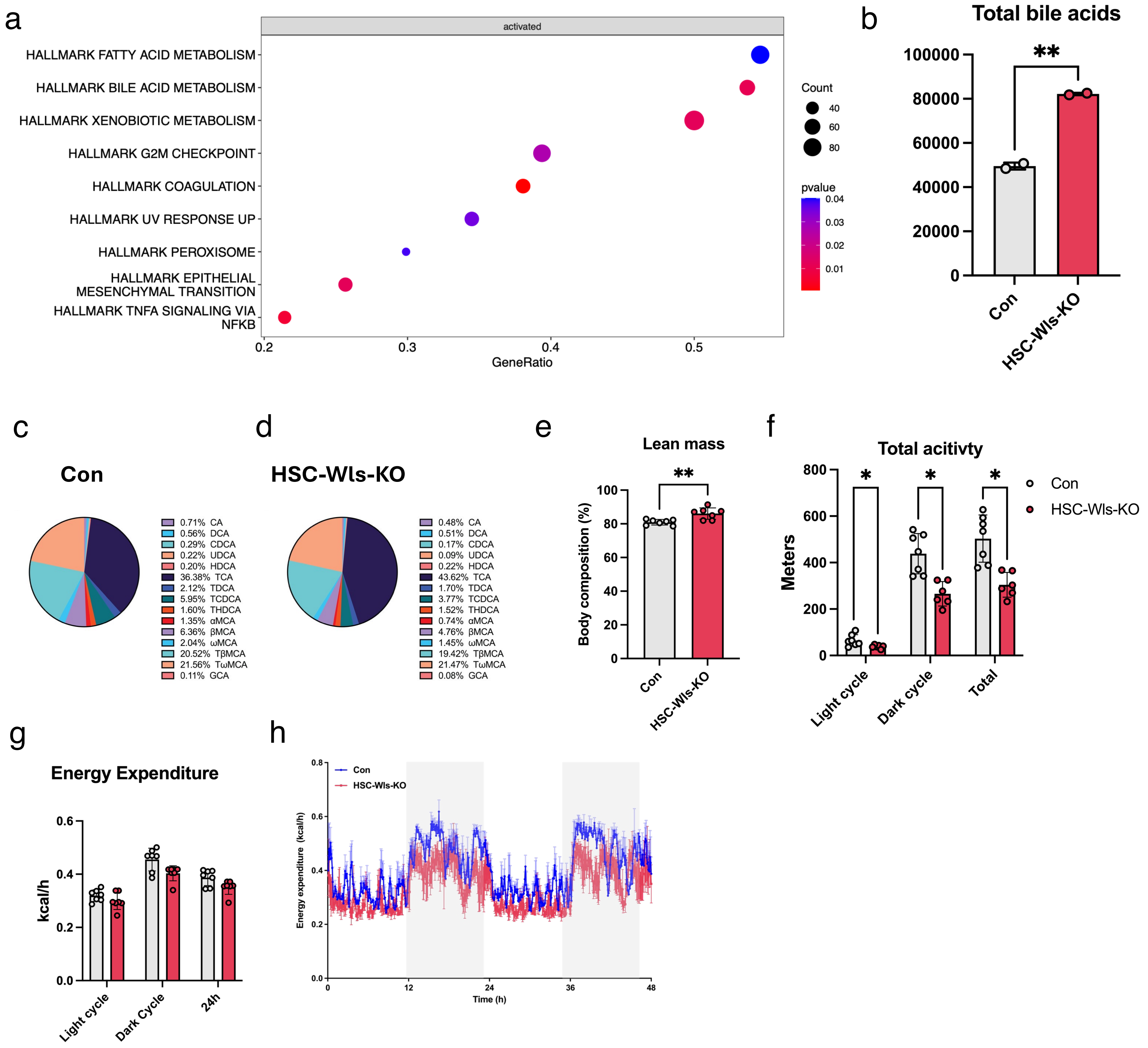


**Figure S9**. **(a)** GSEA plot of enriched hallmark pathways in HSC-Wls-KO versus control hepatocytes using snRNA-seq. **(b)** Total bile acid levels in HSC-Wls-KO versus control livers quantified by mass spectrometry. **(c, d)** Bile acid composition profiles of **(c)** control and **(d)** HSC-Wls-KO livers. **(e)** Lean mass percentage in control and HSC-Wls-KO mice. **(f)** Total locomotor activity (meters) during light cycle, dark cycle, and overall in control and HSC-Wls-KO mice. **(g)** Energy expenditure (kcal/h) during light cycle, dark cycle, and over 24 hours. **(h)** Plot of energy expenditure versus time in HSC-Wls-KO and controls.

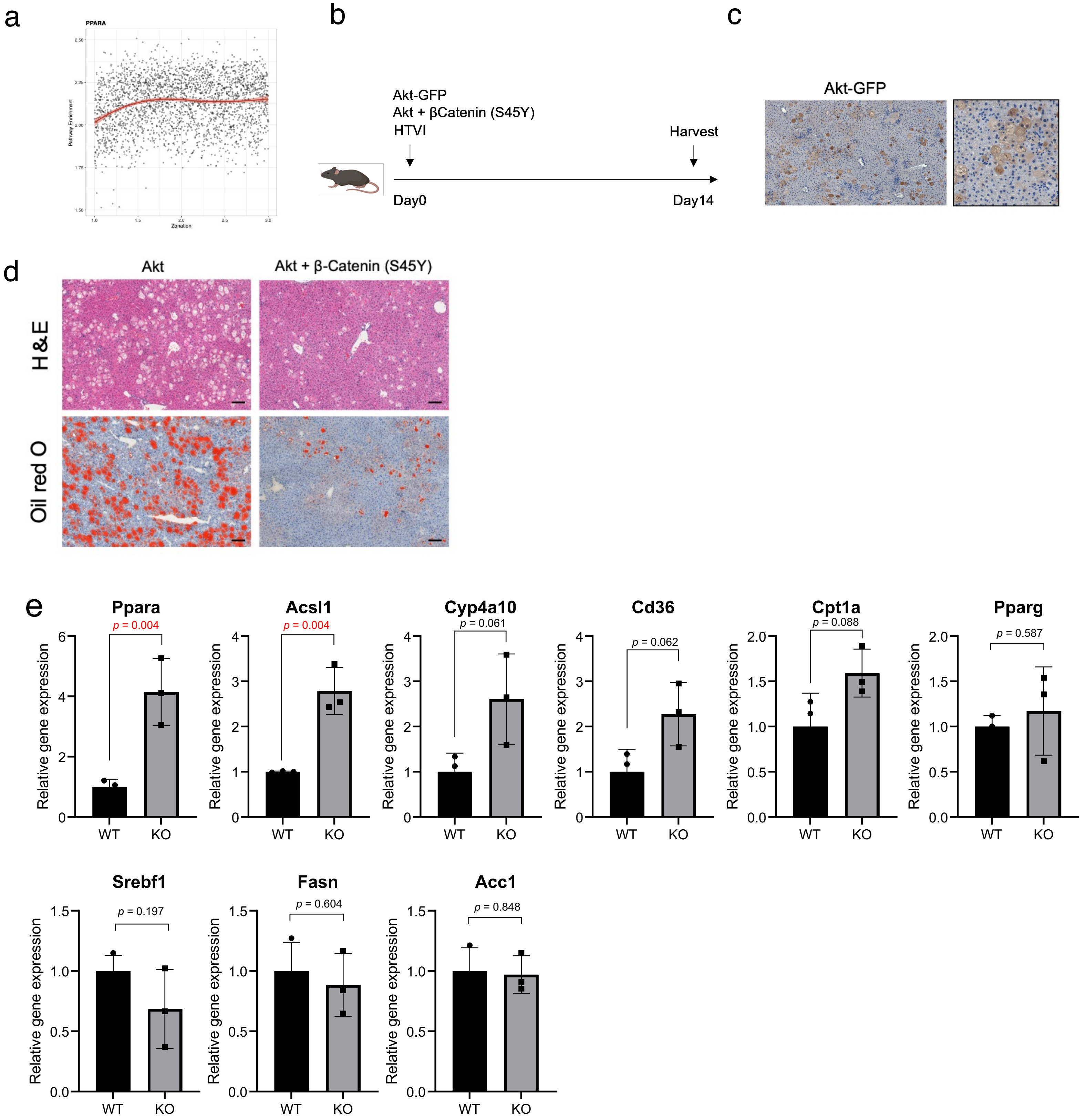


**Figure S10**. **(a)** Scatter plot of PPARα target gene enrichment along the inferred zonation axis in human spatial transcriptomic liver data. The x-axis captures the inferred zonation coordinate as a continuous value from the portal triad (1.0) to the central vein (3.0), while the y-axis indicates the ss-GSEA score. **(b)** Experimental schematic of hydrodynamic tail vein injection (HTVI) of plasmids expressing Akt alone or Akt together with constitutively active β-catenin (S45Y), followed by liver harvest at day 14. **(c)** Representative immunohistochemical staining confirming Akt-GFP expression in hepatocytes following HTVI. **(d)** Histological analysis (H&E and Oil Red O staining) of livers expressing Akt or Akt + β-catenin (S45Y). **(e)** qPCR analysis of genes involved in fatty acid uptake and β-oxidation.

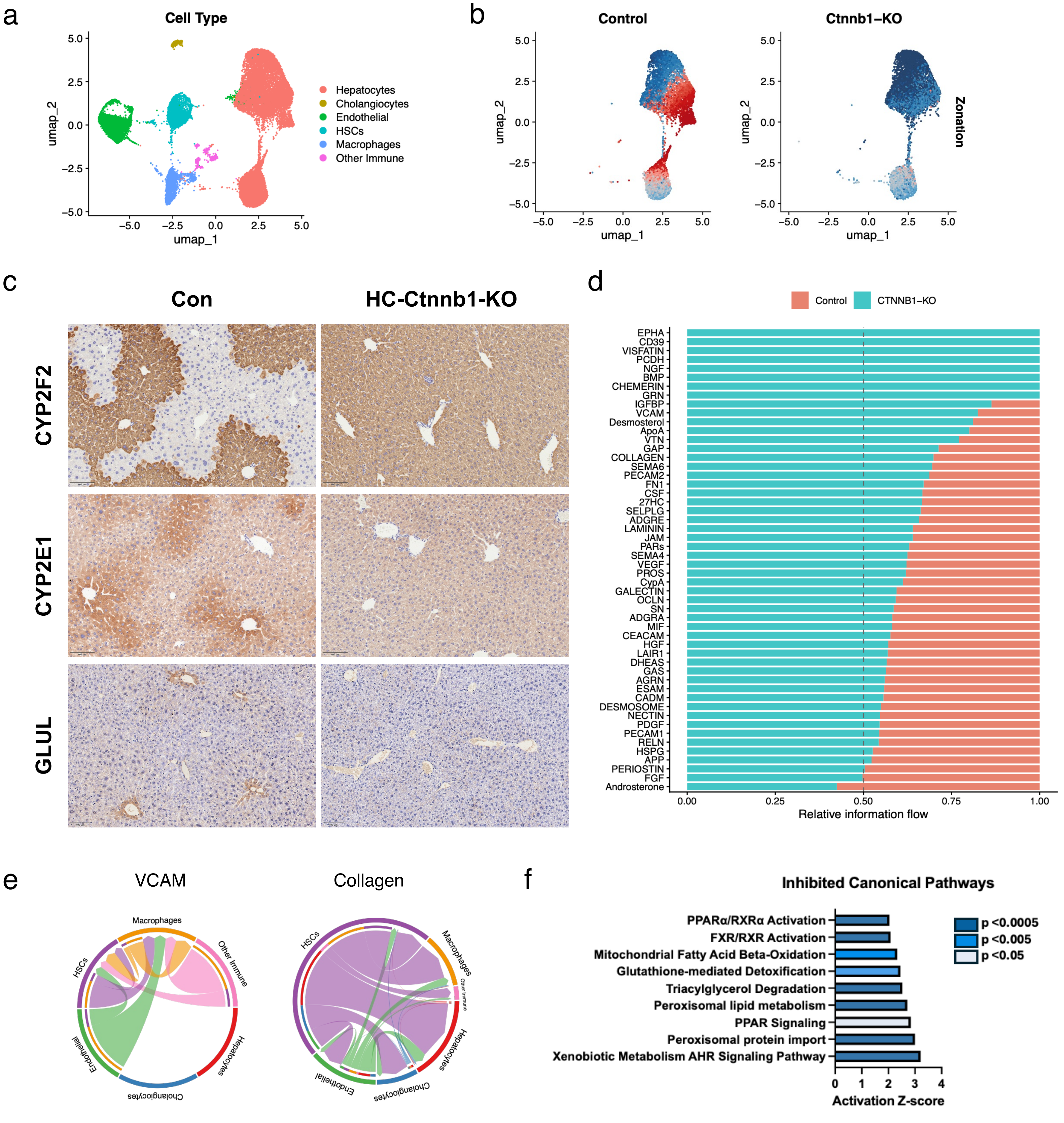


**Figure S11**. **(a)** UMAP plot by cell type in Chromium Flex snRNA-seq of HC-Ctnnb1-KO and control livers. **(b)** UMAP plots of inferred zonal identities using RIZLiver in Chromium Flex snRNA-seq of HC-Ctnnb1-KO and control livers. **(c)** Representative immunohistochemical staining of zonation markers in HC-Ctnnb1-KO and control livers. **(d)** CellChat analysis of relative information flow between Control and HC-Ctnnb1-KO in Chromium Flex snRNA-seq. **(e)** Chord diagrams of VCAM and Collagen cell-cell interaction networks in HC-Ctnnb1-KO livers. **(f)** Inhibited canonical pathways in HC-Ctnnb1-KO mice found by Ingenuity Pathway Analysis (IPA) of Chromium Flex snRNA-seq.

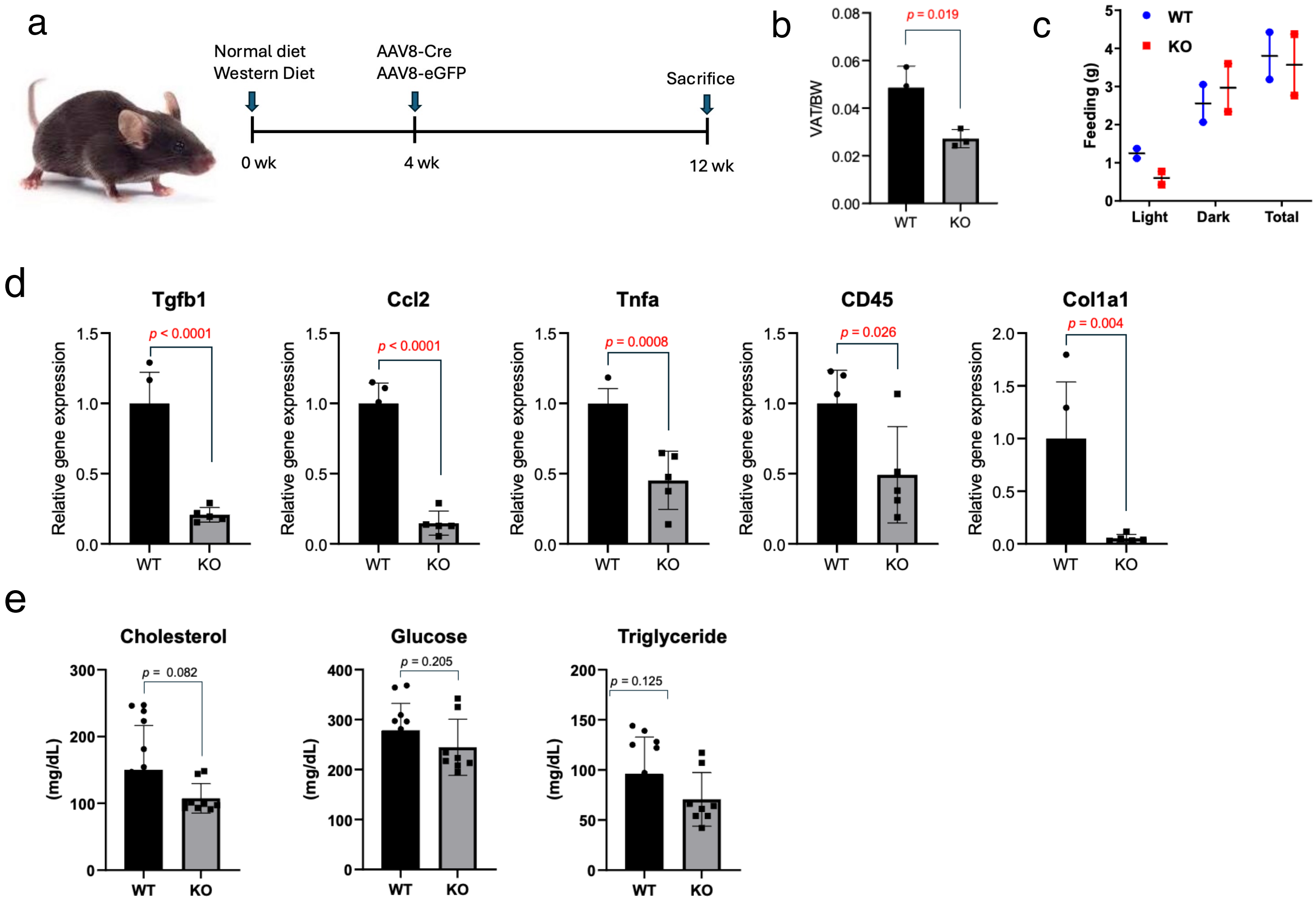


**Figure S12**. **(a)** Experimental schematic comparing normal and western diets in inducible HC-Ctnnb1-KO (using intraperitoneal injection of AAV8-TBG-Cre) versus control mice (AAV8-TBG-GFP) at 4 weeks. Mice were harvested at 12 weeks. **(b)** Quantification of visceral adipose tissue (VAT)-to-body weight ratios (VAT/BW) in HSC-Wls-KO versus controls. **(c)** Food intake during light cycle, dark cycle, and overall. **(d)** qPCR analysis of fibro-inflammatory markers in HSC-Wls-KO versus control livers. **(e)** Serum biochemical analysis of cholesterol, glucose, and triglyceride levels in HSC-Wls-KO versus controls.

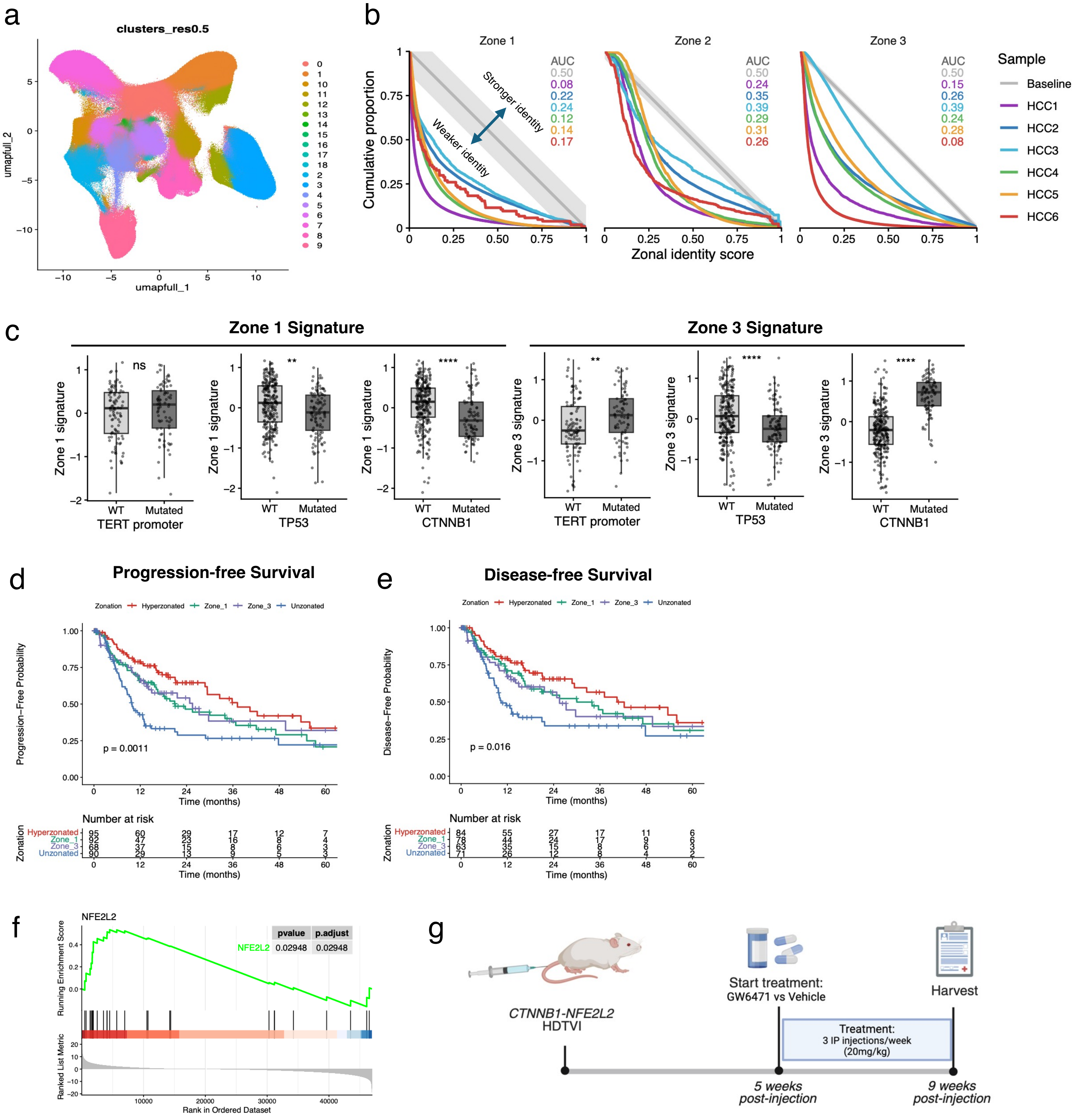


**Figure S13**. **(a)** UMAP plot of Xenium spatial transcriptomics profiling of human control and HCC liver samples. Mice were harvested at 12 weeks. **(b)** Zonal identity scores of cells inferred to be in zones 1, 2 and 3 based on relative lobular position. An AUC above 0.5 indicates overall stronger zonal identity relative to baseline, while an AUC below 0.5 indicates weaker zonal identity relative to baseline. **(c)** Comparison of HCC-specific zone-1 and zone-3 signatures in *TERT* promoter, *TP53*, and *CTNNB1* mutations in TCGA-LIHC HCC tumors. **(d,e)** Kaplan-Meier plots depicting **(d)** progression-free and **(e)** disease-free survival of HCC patients in TCGA-LIHC stratified by zonation subtype. Log-rank p-values as indicated. **(f)** GSEA barcode plot of an *NFE2L2* activation signature in zone-3 tumors in TCGA-LIHC. **(g)** Experimental schematic using the *CTNNB1-NFE2L2* HDTVI model. Mice were treated with a PPARα antagonist, GW6471, via intraperitoneal injection from 5 to 9 weeks post-HDTVI.

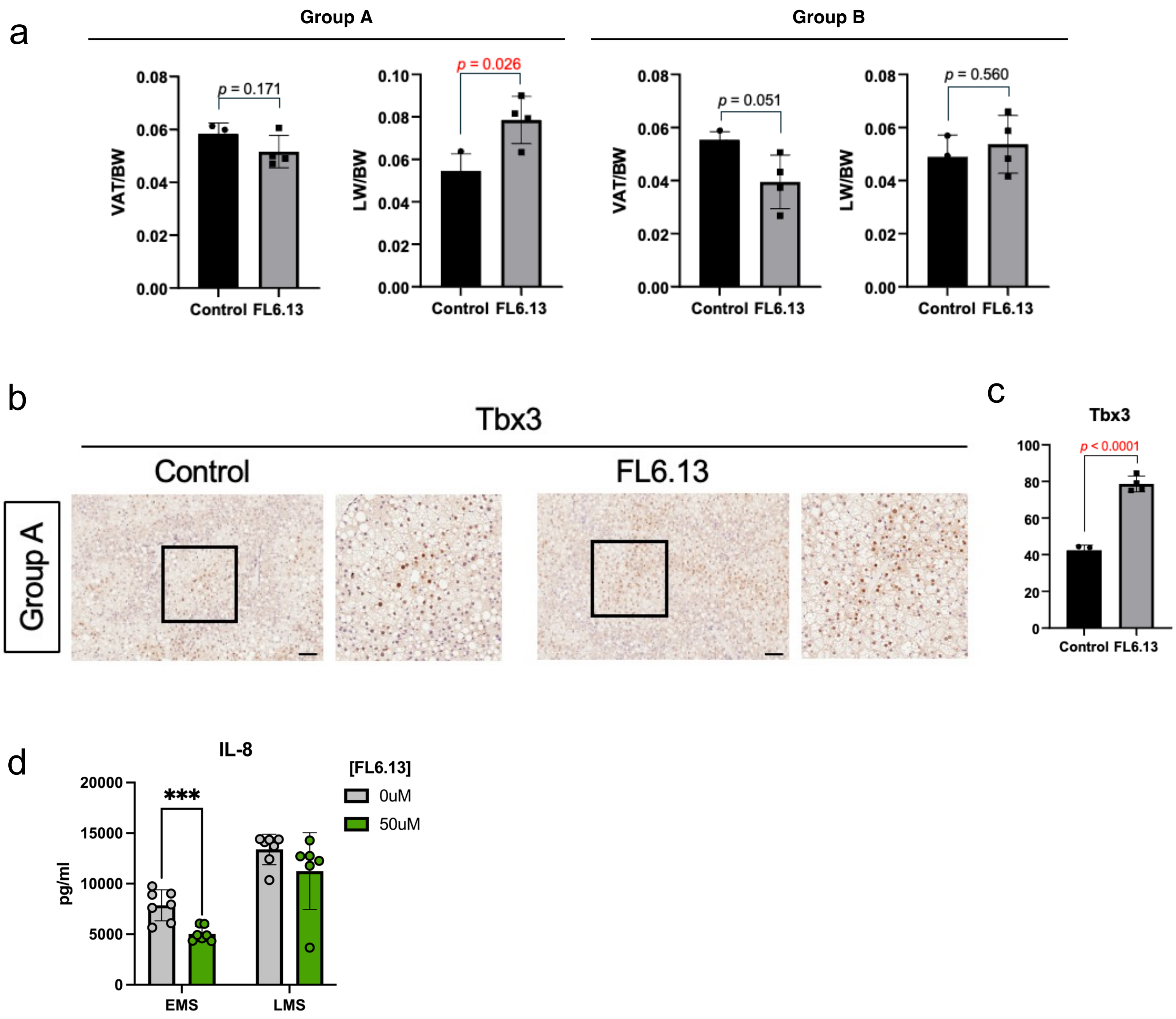


**Figure S14**. **(a)** Quantification of visceral adipose tissue–to–body weight (VAT/BW) and liver weight–to–body weight (LW/BW) ratios in Group A and Group B of FL6.13 treatment in vivo. **(b)** Representative immunohistochemical staining of TBX3 in Group A control and FL6.13-treated livers. **(c)** TBX3 IHC quantification in Group A control and FL6.13-treated livers. **(d)** Cytokine profiling of IL-8 in culture supernatants from early metabolic syndrome (EMS) or late metabolic syndrome (LMS) media, followed by treatment with the Wnt agonist FL6.13 or control.
